# MAP kinase ERK5 modulates cancer cell sensitivity to extrinsic apoptosis induced by death-receptor agonists and Natural Killer cells

**DOI:** 10.1101/2023.03.22.533738

**Authors:** Sergio Espinosa-Gil, Saska Ivanova, Elisenda Alari-Pahissa, Melek Denizli, Beatriz Villafranca-Magdalena, Maria Viñas-Casas, Idoia Bolinaga-Ayala, Andrés Gámez-García, Eva Colas, Miguel Lopez-Botet, Antonio Zorzano, José Miguel Lizcano

## Abstract

Death receptor ligand TRAIL is a promising cancer therapy due to its ability to selectively trigger extrinsic apoptosis in cancer cells. However, TRAIL–based therapies in humans have shown limitations, mainly due inherent or acquired resistance of tumor cells. To address this issue, current efforts are focussed on dissecting the intracellular signaling pathways involved in resistance to TRAIL, to identify strategies that sensitize cancer cells to TRAIL-induced cytotoxicity. In this work, we describe the oncogenic MEK5-ERK5 pathway as a critical regulator of cancer cell resistance to the apoptosis induced by death receptor ligands. Using 2D and 3D cell cultures and transcriptomic analyses, we show that ERK5 controls the proteostasis of TP53INP2, a protein necessary for full activation of caspase-8 activation in response to TNFα, FasL or TRAIL. Mechanistically, ERK5 phosphorylates and induces ubiquitylation and proteasomal degradation of TP53INP2, resulting in cancer cell resistance to TRAIL. Concordantly, ERK5 inhibition or genetic deletion, by stabilizing TP53INP2, sensitizes cancer cells to the apoptosis induced by recombinant TRAIL and TRAIL/FasL expressed by Natural Killer cells. The MEK5-ERK5 pathway regulates cancer cell proliferation and survival, and ERK5 inhibitors have shown anticancer activity in preclinical models of solid tumors. Using endometrial cancer patient-derived xenograft organoids, we propose ERK5 inhibition as an effective strategy to sensitize cancer cells to TRAIL-based therapies and Natural Killer cells.

## Introduction

Apoptosis can be triggered through either the intrinsic or the extrinsic pathways. The intrinsic apoptotic pathway is activated in response to intracellular stimuli that induce cellular damage, and triggers apoptosis through Bcl-2 gene family and activation of caspase-9, which in turn activates the executor caspases -3, -6 and -7 (1). Extrinsic apoptosis is induced by direct interaction of death ligands (TNFα, FasL and TNF-related apoptosis-inducing ligand TRAIL) with their specific death-receptors (DRs) at the plasma membrane. This binding triggers the formation of the death-inducing signaling complex (DISC) comprising FADD, and several molecules of the initiator pro-caspase-8 (2). Then, caspase-8 K63-ubiquitylation by Cullin3 or TRAF6 ligases promotes the aggregation, self-processing and full activation of caspase-8 (3,4). However, efficient caspase-8 K63-ubiquitylation requires of other players to recruit the ligases to the DISC complex. One of these players is TP53INP2, which binds caspase-8 and the TRAF6-Ubc13 dimer in response to DRs activation, facilitating an efficient ubiquitin transfer from Ubc13 to caspase-8 (5).

Among DR ligands TRAIL induces extrinsic apoptosis in a wide variety of cancer cell lines and tumors, without affecting normal cells (6). Therefore, TRAIL-receptor agonists (TRAs) have been extensively investigated as promising cancer therapies, and are currently in clinical trials for several solid cancers (7). However, TRAIL–based therapies in humans have shown limitations, mainly due to their poor ability to activate caspase-8 in solid tumors (7). Hence, to improve TRAIL-based anticancer therapy, it is critical to identify those cellular pathways involved in the sensitization to TRAIL-induced apoptosis.

The MAP kinase ERK5 is activated in response to growth factors and different forms of stress, including oxidative stress and cytokines (8), by direct phosphorylation of its unique upstream kinase MEK5 (9). The MEK5-ERK5 pathway constitutes a unique intracellular axis that mediates cell proliferation and survival of several cancer paradigms, including blood and solid cancers (10,11). Concordantly, elevated levels of ERK5 or MEK5 correlate with bad prognosis and malignancy of several cancers (12–15), and ERK5 inhibitors have shown anticancer activity in tumor xenograft models (16). In addition, ERK5 activity protects cancer cells from entering intrinsic apoptosis, by phosphorylating and inhibiting Bad and Bim pro-apoptotic proteins (17,18).

In this study, we have investigated whether the MEK5-ERK5 pathway also regulates the extrinsic apoptosis induced by DR agonists in cancer cells. We found that ERK5 kinase activity confers resistance to the cytotoxicity exerted by DR agonists in cancer cells, by phosphorylating and inducing proteasomal degradation of TP53INP2. Concordantly, pharmacologic or genetic ERK5 inhibition sensitized cancer cells to apoptosis induced by DR agonists and by TRAIL/FasL expressed by NK cells.

## Results

### ERK5 kinase activity modulates sensitivity of cancer cell to DR agonists

A RNAi-based screening identified ERK5 as one of top 5% genes whose silencing sensitized HeLa cells to TRAIL-induced apoptosis (19). Following studies reported that overexpression of a constitutive nuclear ERK5 mutant favored resistance to TRAIL-induced apoptosis in breast cancer cells (20). These preliminary works prompted us to investigate whether ERK5 pathway is implicated in resistance to TRAIL-induced toxicity in cancer cells.

To investigate the role of ERK5 as a modulator of extrinsic apoptosis, we used endometrial cancer (EC) cells. EC cells frequently show overactivation of PI3K/mTOR (∼50% of EC cells harbour a double allelic mutated PTEN gene) and canonical NF-kB pathways, which protect cancer cells from TRAIL-induced apoptosis (21,22). Thus, EC cells represent a good model of TRAIL-resistant cancer cells. We first investigated the effect of the specific ERK5 inhibitor JWG-071 (14,23) in a panel of human EC cell lines, including type I endometroid (Ishikawa and ANC3A) and type II non-endometroid (ARK1 and ARK2) cells. Cell viability assays showed that pre-treatment with JWG-071 sensitized the four EC cell lines to recombinant TRAIL (rTRAIL)-induced cytotoxicity in a dose-dependent manner (**Fig. 1A**). The structurally different ERK5 inhibitor AX15836 (24) and the MEK5 inhibitors BIX02188 and BIX02189 (25) also sensitized EC cells to TRAIL-induced cell death (**Fig. 1B**). Interestingly, ERK5 inhibition also sensitized EC cells to the toxicity induced by other DR ligands such as TNFα or FasL (**Fig. 1C**), indicating a role for ERK5 kinase activity in mediating the extrinsic apoptosis induced by DR agonists in cancer cells. Finally, we observed that ERK5 inhibition potentiated TRAIL cytotoxicity in a panel of TRAIL-resistant cancer cell lines (**Fig. 1D**), such as neuroblastoma (SK-N-AS), cervical (HeLa), non-small cell lung (A549), and prostate (LnCaP) cancer cells (26–29). Thus, ERK5 inhibition sensitizes cancer cells to TRAIL, independently of cancer type or mutational status.

**Figure 1.**
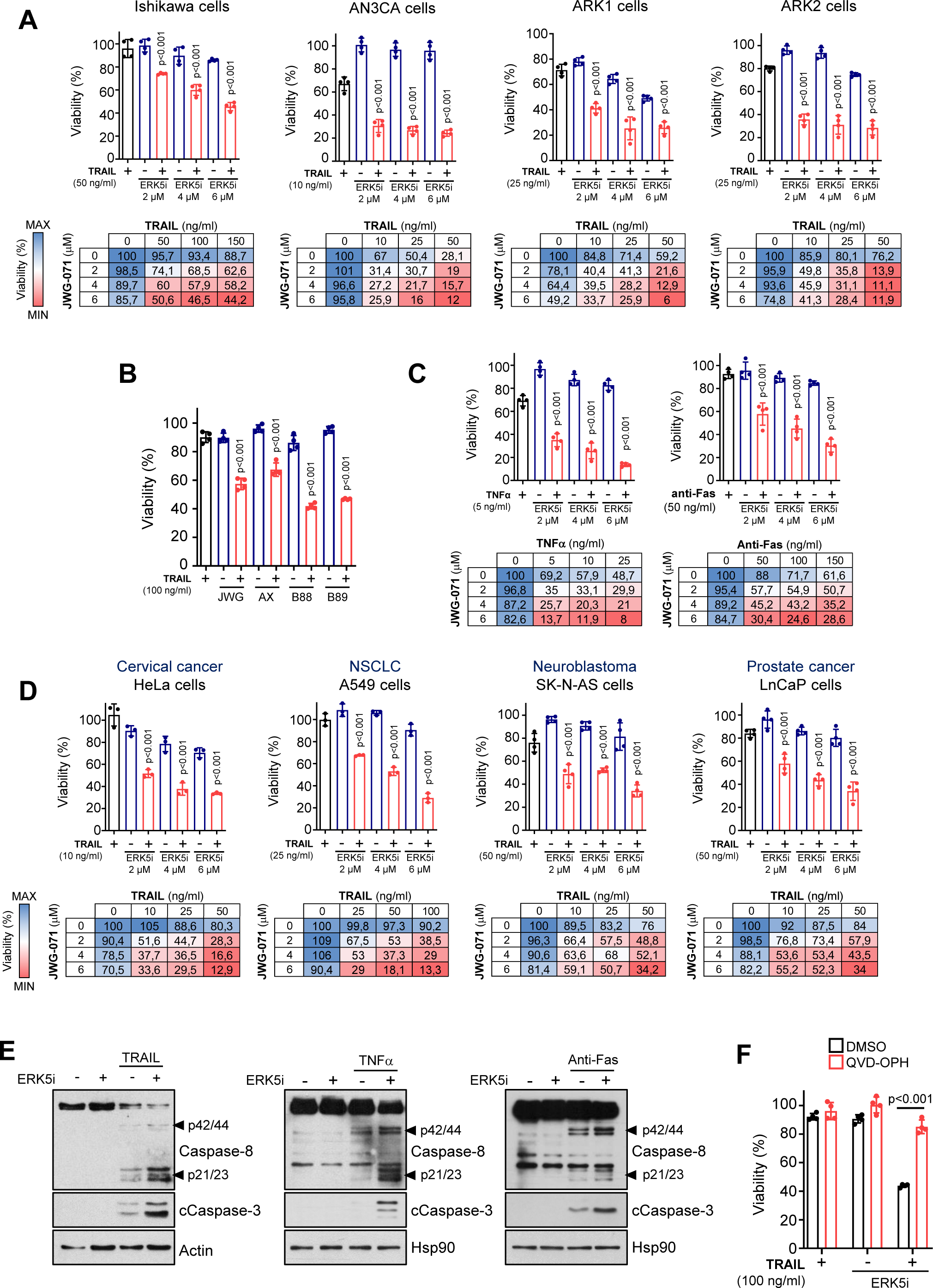
ERK5 inhibition sensitizes cancer cells to DR ligands-induced apoptosis by favoring Caspase-8 activation. **A**, ERK5 inhibition sensitizes EC cells to TRAIL-induced cytotoxicity. EC cells were pre-incubated with ERK5i JWG-071 for 12 h, previous treatment with TRAIL for 24 h (mean ± SD, n = 4 independent biological experiments, one-way ANOVA). Lower tables show cell viabilities (%) of each pairwise combination of drug doses (blue color, maximum cell viability; red color, minimum cell viability). **B,** Inhibitors of the MEK5-ERK5 pathway sensitize Ishikawa EC cells to TRAIL-induced cytotoxicity. EC cells were pre-incubated with different ERK5i (JWG-071 and AX-15836) or MEK5i (BIX02188 and BIX02189) for 12 h, previous treatment with TRAIL for 24 h (mean ± SD, n = 3 independent biological experiments, one-way ANOVA). **C**. ERK5 inhibition sensitizes EC cells to TNF-α and anti-Fas activating antibody. EC cells were pre-incubated with ERK5i JWG-071 for 12 h, previous treatment with TNF-α or anti-Fas for 24 h (mean ± SD, n = 4 independent biological experiments, one-way ANOVA). Lower tables show cell viabilities (%) of each pairwise combination of drug doses (blue color, maximum cell viability; red color, minimum cell viability). **D,** ERK5 inhibition sensitizes different human cancer cell lines to TRAIL-induced cytotoxicity. Cell viability experiments were performed as in **A**. **E,** ERK5 inhibition enhances caspase-8 activation induced by DR ligands. Ishikawa cells pre-treated with 5 μM JWG-071 (12 h) were treated (4 h) with TRAIL, TNFα or anti-Fas activating antibody. Caspase-8 and Caspase-3 activation was monitored by immunoblot analysis. **F,** Inhibition of caspases prevents sensitization to TRAIL induced by ERK5i. Ishikawa cells pre-treated (2 h) with pan-caspase inhibitor Q-VD-OPH were treated with 5 μM JWG-071 (12 h) and further exposed to 100 ng/ml TRAIL for 24h. Histograms show the % of cell viability. Statistical significance was calculated using one-way ANOVA followed by Bonferroni multiple comparison test. Unless indicated, p-values refer to statistical difference of combined (TRAIL and ERK5i) vs TRAIL single treatment.

Ishikawa EC cells exhibited the highest resistance to TRAIL (IC_50_ > 150 ng/ml), and therefore they represent a good model to dissect the mechanism by which ERK5 inhibitors improve TRAIL anticancer activity. Since caspase-8 plays a central role in the extrinsic apoptotic activated by DR ligands(30), we first monitored the activation of caspase-8 and the effector caspase-3 by immunoblot analysis. ERK5 inhibition (ERK5i) potentiated the activation of caspase-8 (and activation of caspase-3) in response to rTRAIL, TNFα or anti-Fas activating antibody, but it had no effect when incubated alone (**Fig. 1E**). In addition, the pan-caspase inhibitor QVD-OPh mostly reverted the cytotoxicity induced by the combined TRAIL and JWG-071 treatment (**Fig. 1F**), demonstrating a role of caspases in ERK5i-mediated sensitization to TRAIL.

The aforementioned results suggest that ERK5 kinase activity protects cancer cells from apoptosis induced by DR ligands. To test this hypothesis, we investigated the impact of the MEK5-ERK5 pathway activity in TRAIL-induced apoptosis in AN3CA EC cells, which were the most sensitive to TRAIL-induced apoptosis (IC_50_, 25 ng/ml). We transiently overexpressed either wild type ERK5 or the ERK5 kinase-dead mutant (ERK5-KD, in which the catalytic loop Asp^200^ was mutated to Ala), in combination with a constitutive active form of MEK5 (MEK5-DD, in which residues Ser^311^ and Ser^315^ were mutated to Asp). Overexpression of active ERK5, but not of kinase dead ERK5, resulted in a significant lower number of apoptotic cells in response to TRAIL treatment, as measured by Annexin V staining and caspase-8 activation assays (**Fig. 2A-B**). Similar results were obtained in Ishikawa EC cells (**Fig. 2C**). Thus, ERK5 kinase activity protects cells from TRAIL-induced apoptosis.

**Figure 2.**
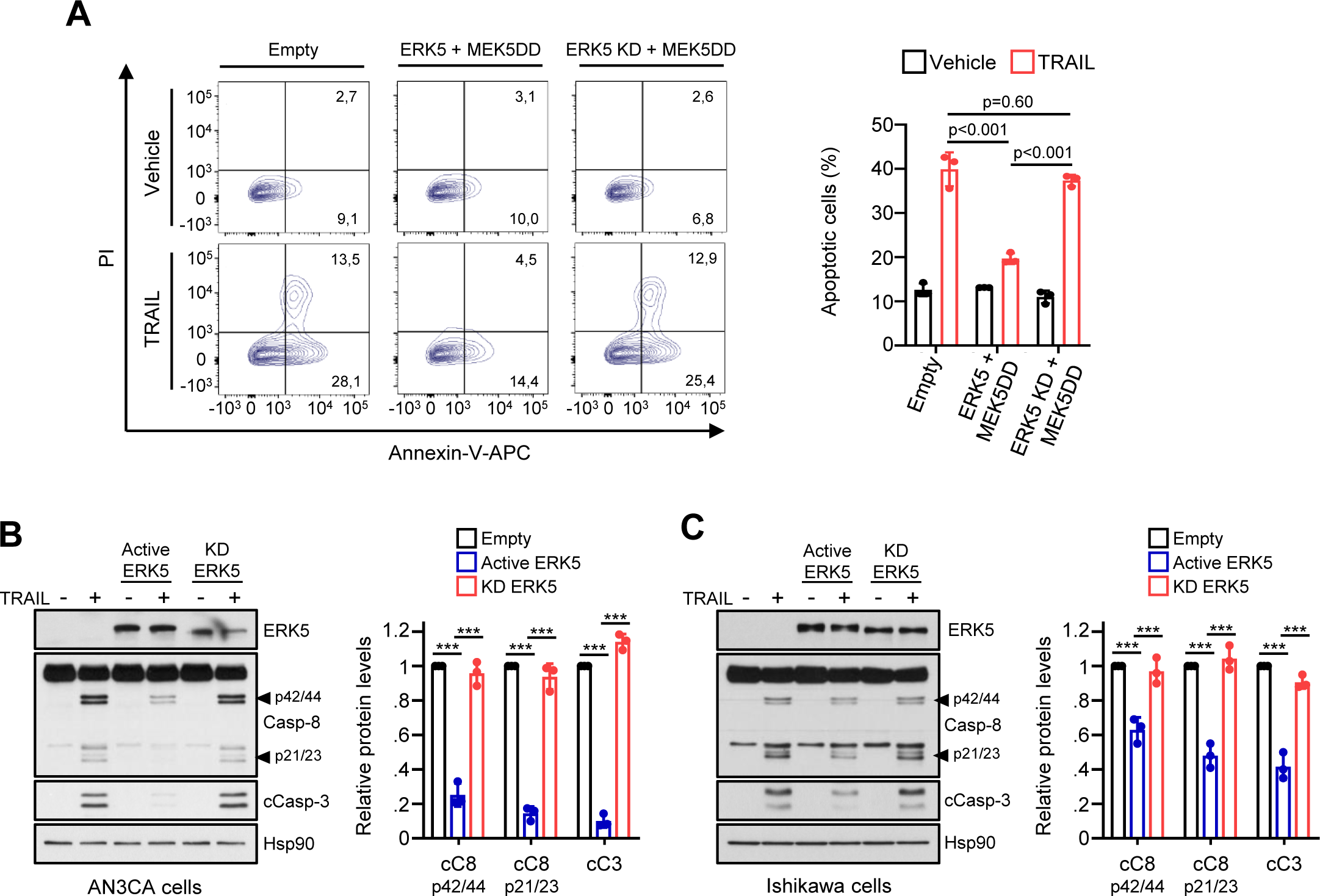
ERK5 kinase activity confers resistance to TRAIL-induced apoptosis in cancer cells. **A**, Active ERK5 protects AN3CA EC cells to TRAIL-induced apoptosis. AN3CA EC cells were transfected with either ERK5 or a ERK5 kinase-dead mutant (D200A, ERK5-KD) together with a constitutively active form of MEK5 (MEK5-DD). Twenty-four hours after transfection cells were treated with 25 ng/ml TRAIL (8 h), and apoptosis was analyzed by flow cytometry (Annexin-V and PI staining). Left panel shows representative results of flow cytometry analysis. Right histograms show the quantification of apoptotic cells (early apoptotic Annexin-V+/PI- and late apoptotic Annexin-V+/PI+ cells). **B-C,** ERK5 kinase activity impairs caspase-8 activation in response to TRAIL in AN3Ca (B) and Ishikawa (C) cells. Immunoblot analysis. Right histograms show the quantification of cleaved caspase-8 and -3 bands (mean ± SD, n = 3 independent biological experiments, one-way ANOVA followed by Bonferroni multiple comparison test). ***p < 0.001.

To obtain genetic evidences on the role of ERK5 activity in sensitization to TRAIL-induced apoptosis, we used CRISPR/Cas9 technology to generate stable Ishikawa cell lines where MEK5 or ERK5 genes were genetically deleted. MEK5 is the only kinase that activates ERK5 by direct phosphorylation of the TEY motif, and ERK5 is the only known substrate described for MEK5 (31). Therefore, MEK5-KO cells lack ERK5 kinase activity. We recently reported that MEK5-KO Ishikawa cells, while lacking ERK5 activity, show normal activation of other MAP kinases (ERK1/2, p38s or JNKs), or components of the PI3K pathway (Akt, mTORC1 or p70S6K) in response to mitogens, stressors or insulin (14). On the other hand, ERK5-KO cells lack both kinase activity-dependent and -independent functions. MEK5 or ERK5 genetic deletion sensitized Ishikawa cells to the extrinsic apoptosis induced by DR agonists TRAIL, TNFα and anti-Fas activating antibody (**Fig. 3A-C**). Similar results were obtained in cervical HeLa and NSCLC A549 MEK5-KO cells, where TRAIL induced higher cytotoxicity compared to wild type cells (**Suppl. Fig. 1**). As seen for the ERK5 inhibitor, ERK5-KO and MEK5-KO Ishikawa cells showed enhance caspase-8 and -3 activation in response to DR ligands (**Fig. 3B and 3C**, respectively). Finally, we tested the cytotoxic effect of DR agonists in 3D cultures of MEK5-WT or MEK5-KO Ishikawa cells, a more complex model to test the effect of drugs. In LIVE/DEAD fluorescence staining assays, 3D cultures of cells lacking ERK5 kinase activity (MEK5-KO cells) showed higher sensitivity in response to TRAIL, TNFα or anti-Fas activating antibody treatment, compared to 3D cultures of wild type cells (**Fig. 3D**).

**Figure 3.**
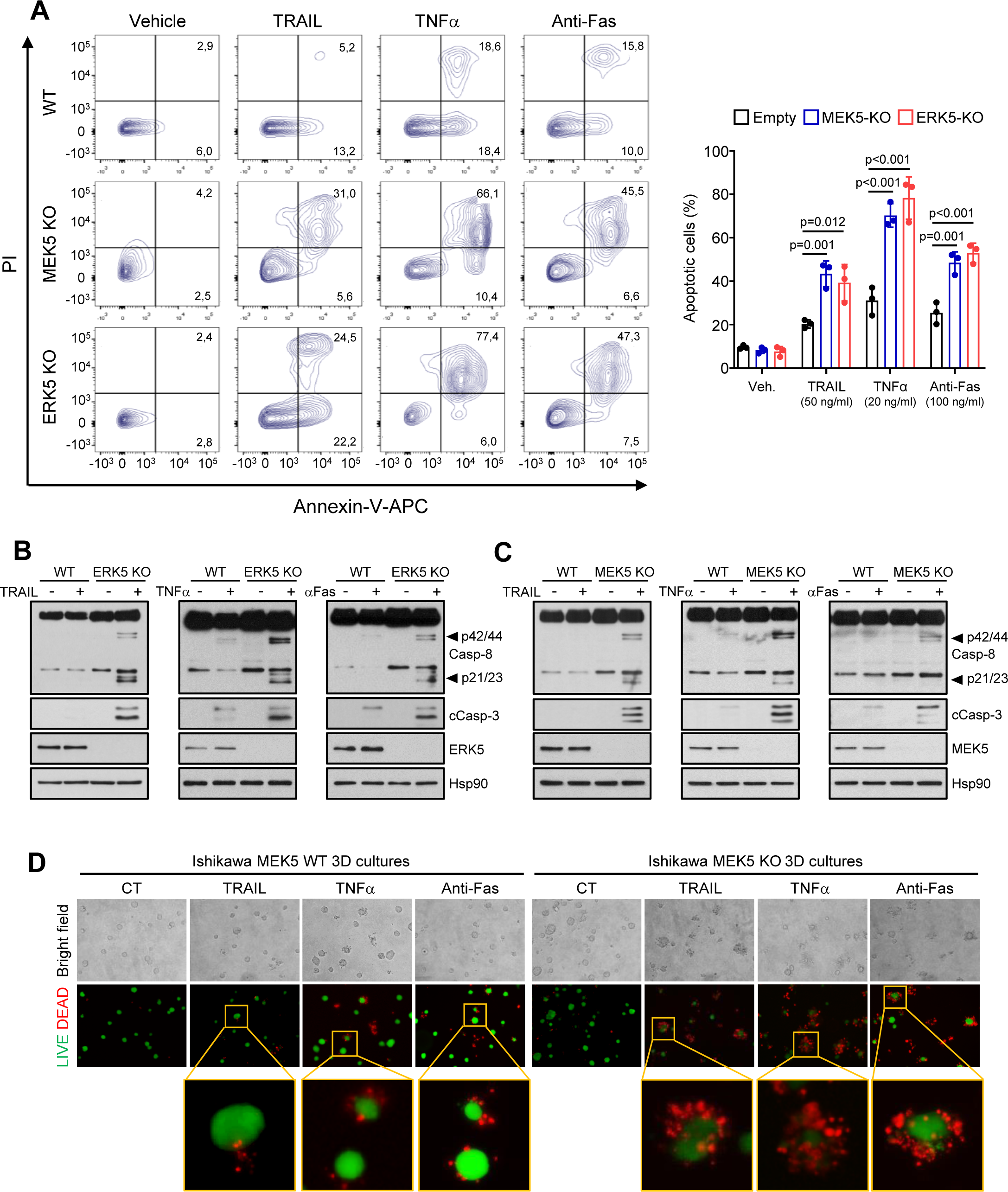
ERK5 or MEK5 genetic deletion sensitizes Ishikawa EC cells to DR agonists-induced apoptosis by favoring Caspase-8 activation. **A**, Ishikawa wild type, MEK5-KO or ERK5-KO cells were treated for 24 h with 50 ng/ml TRAIL, 20 ng/ml TNFα or 100 ng/ml Anti-Fas activating antibody, and apoptotic cells were analyzed by flow cytometry (Annexin-V and PI staining). Right histograms show the quantification of apoptotic cells (mean ± SD, n =3 independent biological experiments, one-way ANOVA). B-C, MEK5 or ERK5 genetic deletion enhances caspase-8 activation in response to DR ligands. Ishikawa wild type, ERK5-KO **(B)** or MEK5-KO **(C)** cells were treated for 4 h with 50 ng/ml TRAIL, 20 ng/ml TNFα or 100 ng/ml Anti-Fas activating antibody. Caspase-8 and caspase-3 activation was monitored by immunoblot analysis. D, MEK5 genetic deletion sensitizes 3D cultures of Ishikawa cells to DR ligands. 3D cultures were treated with 50 ng/ml TRAIL, 20 ng/ml TNFα or 100 ng/ml Anti-Fas activating antibody for 24 h, and cell viability was analyzed with LIVE/DEAD reagent (alive cells, green; dead cells, red). B-C, similar results were obtained in three independent experiments.

Together, our results support a role for ERK5 kinase activity in conferring resistance to TRAIL-induced apoptosis in cancer cells, whereas ERK5 inhibition improves TRAIL anticancer activity by favoring caspase-8 activation.

### ERK5 inhibition sensitizes EC patient-derived xenograft organoids (PDXOs) to TRAIL-induced toxicity

To further explore the therapeutic potential of ERK5 inhibition as a rTRAIL sensitizer agent, we generated EC patient-derived xenograft organoids (PDXOs). EC PDXOs mimic the architecture of the original tissue, reflect the genetic profile of endometrial tumors, and predict patient prognosis (32,33). Thus, PDXOs represent a highly relevant preclinical tool to study endometrial cancer. We generated PDXOs from 2 EC patients (**Suppl. Fig. 2**). Immunofluorescence analysis confirmed the structural organization of the cells into organoids, with the presence of two key glandular epithelial markers, EpCAM (**Fig. 4A**) and E-cadherin, and the mesenchymal and cytoskeleton markers vimentin and F-Actin, respectively (**Fig. 4B**). The positive and heterogeneous staining of the epithelial markers in our PDXOs recapitulated the morphological organization of endometrial tumors (**Fig. 4A-B**). Next, we tested the effect of ERK5i in PDXOs in combination with rTRAIL, by measuring the viability of the PDXOs using two different techniques: intracellular ATP quantification (**Fig. 4C**), and LIVE/DEAD assay (**Fig. 4D**). PDXOs from the two patients showed different sensitivities to TRAIL, confirming that EC PDXOs exhibit patient-specific responses to drugs (32). Thus, TRAIL treatment as a single agent decreased the viability of PDXOs from patient 440, but did not affect viability of PDXOs from patient 1297 even at high concentrations (50 ng/ml). Interestingly, pre-incubation with the ERK5i sensitized both PDXOs to TRAIL-cytotoxicity (**Fig. 4B-C**). Together, our results suggest ERK5i as a promising strategy to sensitize EC tumors to TRAIL anticancer activity, independently of the sensitivity status of the patients to TRAIL.

**Figure 4.**
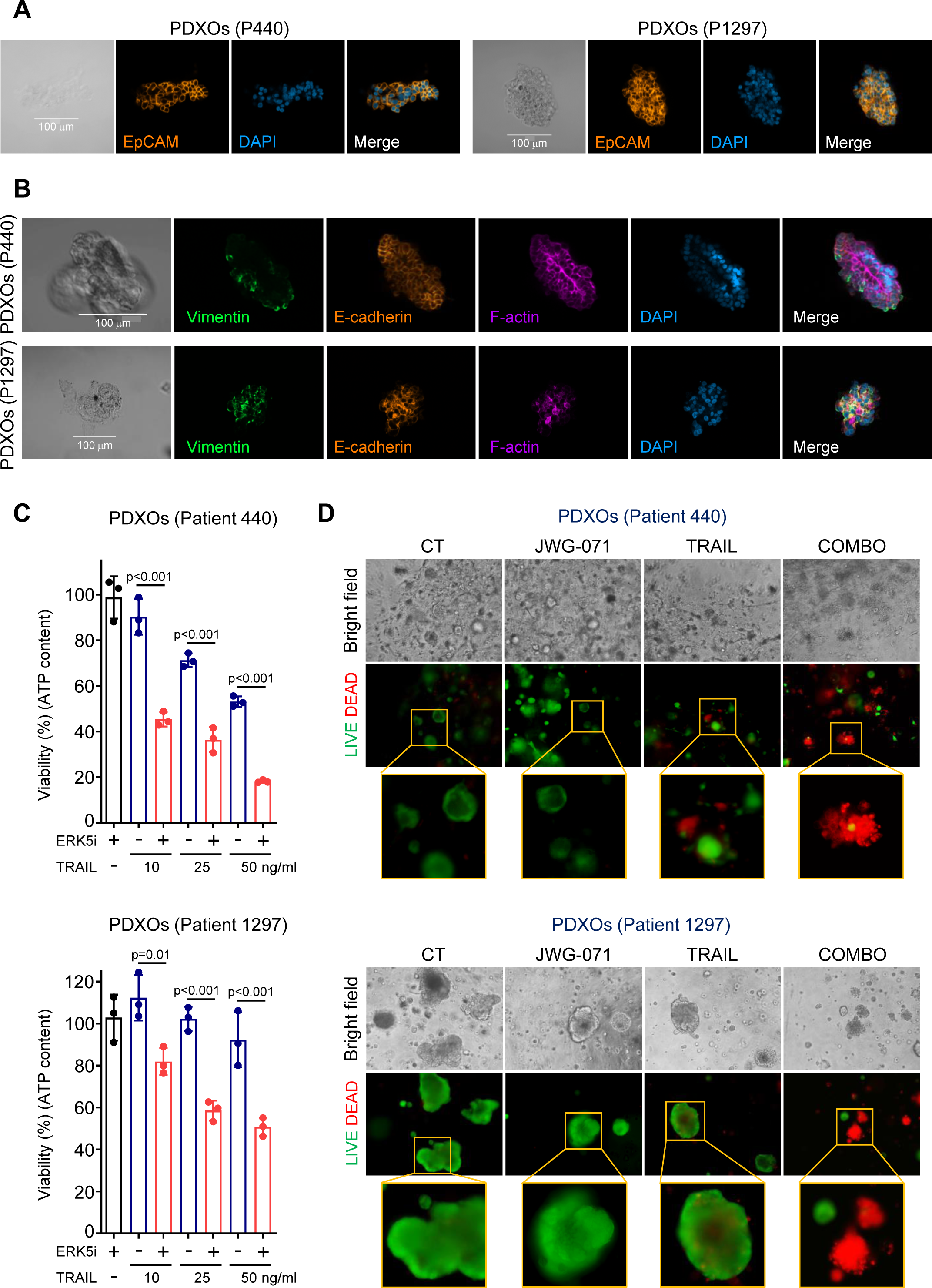
ERK5 inhibition sensitizes PDX-derived organoids (PDX-Os) from endometroid cancer patients to TRAIL-induced cytotoxicity. **A-B**, PDX-Os recapitulates the morphological organization of endometrial tumors. Confocal immunofluorescence imaging of PDX-Os from patients 440 and 1297 stained with epithelial (EpCAM and E-cadherin), mesenchymal (vimentin), cytoskeleton (F-actin) and nuclei (DAPI) markers. Scale bars are represented in the bright field photos. **C-D**, ERK5 inhibition sensitize EC PDX-Os to TRAIL-induced cytotoxicity. PDX-Os from patients 440 and 1297 were pre-treated with either vehicle or 5 µM JWG-071 for 12 h, previous treatment with TRAIL (48 h). Cytotoxicity was assessed by measuring intracellular ATP levels in PDXOs **(C)**, or by LIVE/DEAD staining of PDX-Os **(D)** (alive cells, green; dead cells, red). Statistical significance was calculated using one-way ANOVA followed by Bonferroni multiple comparison test. Individual p-values are indicated in each figure.

### ERK5 inhibition or genetic deletion induces TP53INP2 protein levels to sensitize cancer cells to DR agonists

TP53INP2 mediates caspase-8 K63-ubiquitylation in response to DR agonists, facilitating caspase-8 activation and apoptosis induced by these agonists (5). Hence, we next investigated the effect of ERK5 modulation on TP53INP2 protein levels in four EC cell lines. EC cells treated with JWG-071 showed a robust increase on TP53INP2 protein levels (**Fig. 5A**), without affecting DR5 protein levels. Furthermore, ERK5 or MEK5 genetic deletion also resulted in increased TP53INP2 protein levels (**Fig. 5B**). On the other hand, caspase-8 full activation in response to DRs activation requires the binding of TP53INP2(5). Interestingly, in cells transiently overexpressing FLAG-tagged TP53INP2, ERK5 inhibition resulted in higher levels of TP53INP2/Caspase-8 complex (**Fig. 5C**).

**Figure 5.**
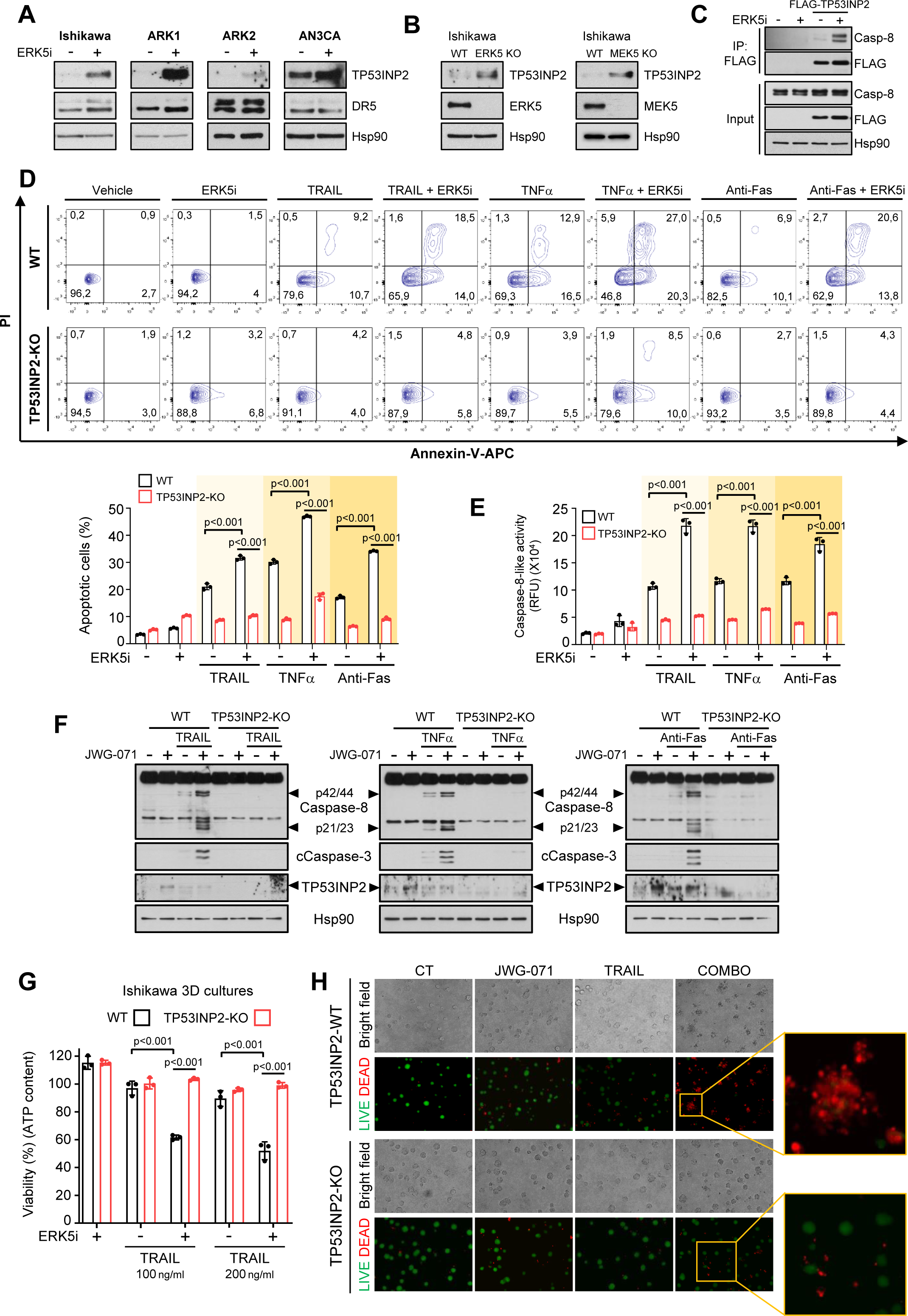
ERK5 inhibition or genetic deletion sensitize cancer cells to DR agonists by increasing TP53INP2 protein levels. **A**, ERK5 inhibition upregulates TP53INP2 protein levels in EC cancer cells. Ishikawa, ARK1, ARK2 and AN3CA EC cells were treated for 24 h with either vehicle or 5 μM JWG-071. TP53INP2 and DR5 protein levels were determined by immunoblot analysis. **B,** ERK5 or MEK5 genetic deletion upregulates TP53INP2 protein levels. **C,** ERK5 inhibition increases the amount of TP53INP2 bound to Caspase-8. HEK293T cells overexpressing FLAG-TP53INP2 were treated with either vehicle or 3 μM JWG-071 for 6 h. After immunoprecipitation of FLAG-TP53INP2, the presence of Caspase-8 in the immunoprecipitated fraction was detected by immunoblot analysis. **D**. TP53INP2 mediates the sensitization of EC cells to DR agonists exerted by ERK5i. Ishikawa wild type TP53INP2-KO cells were pre-treated 3 μM JWG-071 (16 h), treated with 50 ng/ml TRAIL, 20 ng/ml TNFα or 100 ng/ml Anti-Fas activating antibody for 24 h, and apoptotic cells were analyzed by flow cytometry (Annexin-V and PI staining). Lower panel shows the corresponding quantification of the results of three independent biological experiments (one-way ANOVA followed by Bonferroni multiple comparison test). **E-F,** ERK5 inhibition favors caspase-8 activation in response to DR-ligands in a TP53INP2 protein-dependent manner. Caspase-8 activation was measured by a luminescent substrate (LETD) assay (**E**), or by monitoring Caspase-8 cleavage by immunoblot (**F**). Cells were pre-treated with JWG-071 (12 h) and further treated with 50 ng/ml TRAIL, 20 ng/ml TNFα or 100 ng/ml Anti-Fas activating antibody for 4 h. **G-H,** ERK5 inhibition sensitize 3D cultures of EC Ishikawa cells to TRAIL in a TP53INP2-dependent manner. 3D cultures of Ishikawa wild type or TP53INP2-KO cells were pre-treated 12h with JWG-071, and further treated with the indicated concentrations of TRAIL for 24 h. Viability of the 3D cultures was assessed by measuring intracellular ATP-content (**G**) or by LIVE/DEAD staining (**H**). Alive (green) and dead cells (red) were visualized by fluorescence microscopy. Statistical significance of **E** and **G** was calculated using one-way ANOVA followed by Bonferroni multiple comparison test. Individual p-values are indicated in each panel.

Next, we investigated whether TP53INP2 mediates the sensitization to DR ligands exerted by ERK5 inhibition. We generated CRSIPR/Cas9 TP53INP2-KO Ishikawa cells, to assess the apoptosis induced by DR agonists after pre-treatment with the ERK5i. Compared to wild type Ishikawa cells, TP53INP2-KO cells showed lower apoptosis in response to TRAIL, TNFα or FasL (**Fig. 5D**), confirming TP53INP2 as a player of the extrinsic apoptosis pathway (5). Importantly, EC cells sensitization to DR agonists induced by ERK5 inhibition was greatly impaired in cells lacking *TP53INP2* (**Fig. 5D**). These results were further confirmed in cervical cancer HeLa cells where TP53INP2 was knocked-down (**Fig. Suppl. 3)**. Accordingly, in EC TP53INP2-KO cells, ERK5 inhibition did not enhance caspase-8 activity induced by DR agonists (**Fig. 5E-F**). Finally, ERK5i sensitized Ishikawa-WT 3D cultures to TRAIL-induced toxicity, but not TP53INP2-KO 3D cultures (**Fig. 5G-H**). These results indicate that TP53INP2 mediates the sensitization to DR agonists exerted by ERK5 inhibition, by favoring caspase-8 activation.

### ERK5 phosphorylates and modulates TP53INP2 protein stability

We next investigated whether ERK5 activity could regulate TP53INP2 protein stability, a protein with a short half-life (4 h, (34)). We first studied the role of the proteasome in TP53INP2 proteostasis using cells ectopically expressing human FLAG-TP53INP2 (to avoid an effect due to transcriptional regulation). Short-time treatment (6 h) with the proteasome inhibitor MG-132 resulted in a robust increase on FLAG-TP53INP2 protein levels (**Fig. 6A**). Of note, active ERK5 overexpression - but not ERK5 kinase-dead overexpression-resulted in impaired TP53INP2 protein levels, which were recovered by proteasome inhibition (**Fig. 6A**). We discarded a direct effect of MEK5 or ERK5 on the proteasome activity, since MEK5 or ERK5 inhibitors did not induce accumulation of cellular ubiquitylated proteins **(Fig. Suppl. 4).** These results suggested that ERK5 kinase activity mediates the TP53INP2 ubiquitylation required for its proteasomal degradation. This was confirmed by denaturing pull-down assays using cells overexpressing His-tagged ubiquitin, TP53INP2 and/or active ERK5. Proteasome inhibition increased TP53INP2 protein levels and accumulation of ubiquitylated species (**Fig. 6B**). Importantly, overexpression of active ERK5 induced TP53INP2 ubiquitylation and proteosomal degradation, but not of the ubiquitylation-deficient TP53INP2-3K/R mutant (35) (**Fig. 6C**). On the contrary, ERK5 inhibition restored the low TP53INP2 protein levels induced by the protein synthesis inhibitor cycloheximide (**Fig. 6D**, **upper panels)**. As a control, cycloheximide treatment did not affect protein expression of the ubiquitylation-deficient TP53INP2-3K/R mutant (**Fig. 6D**, **lower panels).**

**Figure 6.**
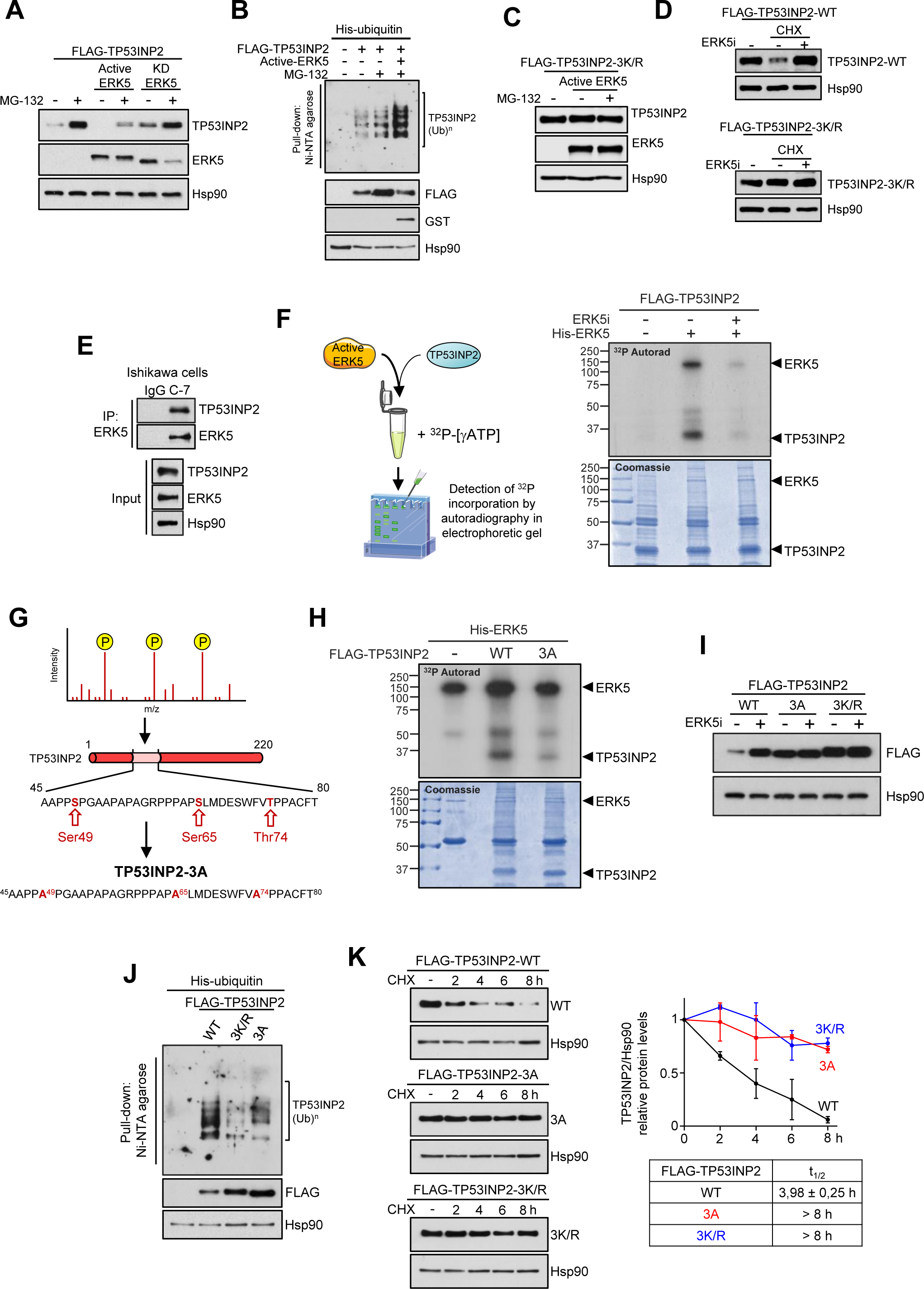
ERK5 phosphorylation of TP53INP2 induces ubiquitylation and proteasomal degradation of TP53INP2. **A**, Active ERK5 overexpression induces TP53INP2 proteasomal degradation. HEK293T were transfected with vectors encoding for FLAG-TP53INP2 and GST-tagged wild type ERK5 (ERK5-WT) or ERK5 kinase-dead mutant (ERK5-KD), in combination with a vector encoding for a constitutively active form of MEK5 (MEK5-DD). Cells were treated with either vehicle or MG-132 (4 h), and protein levels monitored by immunoblot analysis. **B**. ERK5 kinase activity induces TP53INP2 ubiquitylation. HEK293T cells were transfected with plasmids encoding for FLAG-TP53INP2 and His-tagged ubiquitin, and/or for GST-tagged ERK5 and MEK5DD (active ERK5). After incubating with MG-132 (4 h), ubiquitylated TP53INP2 was affinity-purified and detected by immunoblotting. **C,** Active ERK5 does not induce proteasomal degradation of the ubiquitin-deficient mutant TP53INP2-3K/R. Experiment was performed as in **A**. **D,** ERK5 inhibition stabilizes TP53INP2 protein levels. Immunoblot analysis of HEK293 cell lysates overexpressing wild type or ubiquitin-deficient mutant, treated with ERK5i and the protein synthesis inhibitor cyclohexamide (CHX) for 8 hours. **E,** ERK5 interacts with TP53INP2. One milligram of Ishikawa cell lysates was immunoprecipitated with anti-ERK5 antibody, and levels of pelleted TP53INP2 and ERK5 were monitored by immunoblot analysis. **F-G**, ERK5 phosphorylates TP53INP2 at Ser49, Ser65 and Thr74. **F,** FLAG-TP53INP2 protein expressed in HEK293T was affinity purified, and incubated with pure active ERK5 in the presence of [γ-^32^P]-ATP. After resolving the proteins by SDS/PAGE electrophoresis, incorporation of ^32^P was monitored by autoradiography. **G,** Schematic representation of the TP53INP2-3A phospho-deficient mutant. **H,** The TP53INP2-3A mutant is poorly phosphorylated by ERK5. Experiment was performed as in **F**. **I,** The TP53INP2 mutants TP53INP2-3A and TP53INP2-3K/R have higher protein expression than wild-type TP53INP2, and ERK5i does not affect protein levels of TP53INP2-3A and TP53INP2-3K/R mutants. Immunoblot analysis. **J-K**, ERK5 phosphorylation induces TP53INP2 ubiquitylation and proteasomal degradation. **J**, Denaturing ubiquitylation assay of HEK293T cell lysates. Experiment was performed as described in **B**. **K,** Immunoblot analysis of cells expressing wild-type, 3A or 3K/R mutants of TP53INP2, after blocking protein synthesis with cyclohexamide for the indicated times. Right panel shows the quantification of three independent experiments, and the half-life values for each of the TP53INP2 mutants.

Phosphorylation of KLF2 and cMYC by ERK5 regulates ubiquitylation and the proteostasis of these transcription factors (36,37). Given our results, we next explored whether this was also the case for TP53INP2. As it happens for other MAPKs, ERK5 requires to dock the substrate for an efficient phosphorylation. Immunoprecipitation assays using cell lysates with endogenous (**Fig. 6E**) or overexpressed GST-tagged ERK5 and FLAG-tagged TP53INP2 proteins (Fig. Suppl. 5), showed that both proteins interact with each other. Next, we performed in vitro radiochemical kinase assays, incubating affinity-purified TP53INP2 with pure recombinant active ERK5 (38) in the presence of ^32^P-[γATP]. ERK5 efficiently phosphorylated TP53INP2 in vitro (**Fig. 6F**). TP53INP2 phosphorylation and ERK5 autophosphorylation were ablated with the ERK5 inhibitor JWG-071 (**Fig. 6F**), indicating that trace contaminant kinases were not responsible for the phosphorylation.

To identify the TP53INP2 residues phosphorylated by ERK5, phosphorylated TP53INP2 was subjected to tryptic digestion and LC-MS/MS analysis. However, TP53INP2 is a proline-rich protein with low content of basic residues, and tryptic digestion only allowed the identification of peptides comprising the N-term (aa 1-23) and C-term (aa 145-220), where Thr12, Ser14 and Ser208 were identified equally phosphorylated in both control and ERK5 samples (**Suppl. Fig. 6**). These results suggested that ERK5 phosphorylates TP53INP2 central region (aa24-144). ERK5 is a proline-directed kinase that phosphorylates Ser/Thr residues immediately preceded/followed by a proline residue (39). TP53INP2 central region contains two clusters of Ser/Thr residues preceded or followed by a proline: cluster 1, containing Ser49/Ser65/Thr74; and cluster 2, containing Ser94/Ser105/Ser121/Ser123/Thr141. We generated a TP53INP2 mutant in which the three putative phosphorylated sites in cluster-1 were mutated to Ala (TP53INP2-S49A/S65A/T74A, hereafter called TP53INP2-3A) (**Fig. 6G**). This mutant retained the ability to interact with ERK5 (**Suppl. Fig. 7**), but it was poorly phosphorylated by ERK5 (**Fig. 6H**), indicating that ERK5 at least phosphorylates Ser49, Ser65 and Thr74 in TP53INP2.

Next, we sought to investigate the impact of ERK5 phosphorylation in TP53INP2 stability, by comparing the ectopic expression in cells of wild type, phospho-deficient and ubiquitylation-deficient mutants. Both phospho-deficient and ubiquitylation-deficient mutants showed similar expression, which was higher than wild type TP53INP2 (**Fig**. **6I**). Furthermore, ERK5 inhibition increased the levels of wild-type TP53INP2 protein, but not of any of the mutants. Of note, ERK5 inhibition resulted in wild type TP53INP2 protein levels similar to those observed for basal expression of the 3A or 3K/R mutants (**Fig. 6I**). These results prompted us to investigate whether ERK5 phosphorylation affected TP53INP2 ubiquitylation, in denaturing pull-down ubiquitin experiments. Both ubiquitylation-deficient and phospho-deficient TP53INP2 mutants showed impaired ubiquitylation, compared to the wild type protein, suggesting impaired proteasomal degradation (**Fig. 6J**). This was confirmed in cycloheximide chase experiments: ectopically expressed TP53INP2 showed a half-life of ∼4 hours, while the phospho-deficient-3A mutant behaved as a more stable protein (similar to the ubiquitylation-deficient mutant, half-life > 8 hours) (**Fig. 6K**). Together, our results suggest that phosphorylation at Ser49, Ser65 and Thr74 by ERK5 induces TP53INP2 ubiquitylation and proteasomal degradation.

### ERK5 inhibition does not affect apoptosis-related transcriptional programs

To investigate changes on transcriptional programmes related to apoptosis due to ERK5 inhibition, we performed RNA-sequencing of EC Ishikawa cells treated with JWG-071. Only 260 genes were differentially expressed (DEG) in response to ERK5 inhibition (Qvalue<0.05, log_2_ fold change >|1.5|) (**Fig. 7A**), of which 202 (∼80% of DEGs) were downregulated (**Figure 7B**), suggesting that ERK5 kinase activity positively regulates transcription in EC cells. Altered hallmarks include angiogenesis, p53-pathway and cholesterol hallmarks, but not the apoptosis hallmark (**Fig. 7C**). Gene Set Enrichment Analysis (GSEA) did not identify significant alterations (FDR=0.44) of global apoptosis or extrinsic apoptosis (FRD=0.51) transcriptional programs in response to ERK5i (**Fig. 7D-F**). Interestingly, ERK5i did not alter *TP53INP2* gene transcription, supporting a posttranslational regulation of TP53INP2 protein by ERK5.

**Figure 7.**
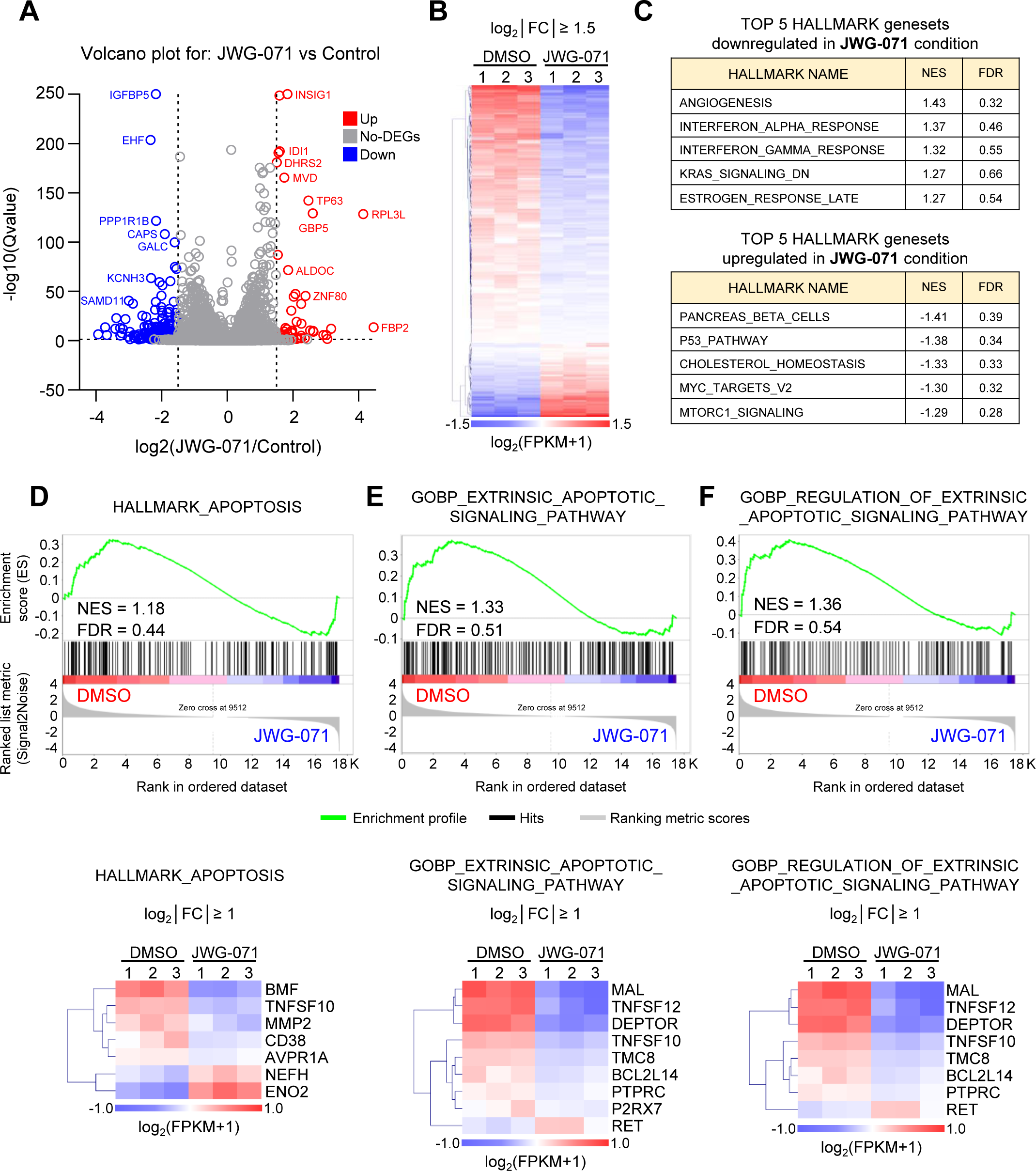
ERK5 inhibition does not alter apoptotic transcriptional programs in Ishikawa EC cells. **A.** Volcano plot of RNAseq expression data after ERK5 inhibition in EC Ishikawa cells. The plot shows the upregulated (in red) and downregulated (in blue) differentially expressed genes (DEGs) (Qvalue < 0.05, log_2_ fold change >|1.5|). Source data is provided as a Supplementary Data file. **B,** ERK5 kinase activity positively regulates gene transcription in EC cells. Heatmap of the 260 DEGs (Qvalue < 0.05, log_2_ fold change >|1.5|) in response to ERK5 inhibition. **C,** Top-5 upregulated and top-5 downregulated hallmark genesets in response to ERK5 inhibition. **D-E-F** Gene set enrichment analysis (GSEA) of Hallmark_apoptosis (**D**), or GOBP-extrinsic apoptotic signaling pathway **(E**) or GOBP-regulation of extrinsic apoptotic signaling pathway (**F**), in response to ERK5 inhibition. Lower panels show the corresponding heatmaps of the DEGs (Q value < 0.05, log_2_ fold change >|1|).

Overall, transcriptomic analysis supports the hypothesis that ERK5 modulates extrinsic apoptosis by posttranslational mechanisms, rather than altering transcription of apoptosis-related genes, at least in EC cells.

### ERK5 inhibition or its genetic deletion sensitize EC cells to TRAIL/FasL expressed by Natural Killer cells

Natural killer cells induce intrinsic and extrinsic apoptosis in cancer cells by two different mechanisms. Intrinsic apoptosis results from the action of the cytolytic pathway, induced by the release of perforin and granzymes that directly cleave and activate caspase-3. On the other hand, NK cells express membrane TRAIL and FasL, which bind DR4/5 (TRAIL receptors) and Fas (CD95) in target cells, triggering extrinsic apoptosis in tumor cells (40).

We studied whether ERK5 inhibition sensitizes EC cells to apoptosis induced by NK cells. We expanded NK cells from freshly isolated PBMCs of four healthy human donors and purified them. eNK cells displayed phenotypic differences compared to fresh NK cells (**Suppl.** **Fig 8**), and expressed higher levels of activating receptors involved in the response to tumor cells (e.g. NKG2D, NKp46, NKp30) (41) (**Suppl. Fig 9 A-C**). eNK cells expressed granzyme B and perforin (**Supp. Fig. 9 D and E**) and displayed surface TRAIL (**Fig. 8A**) and FasL, the latter detected only following incubation with tumor cells (**Fig. 8B**), in agreement with a previous report (40).

**Figure 8.**
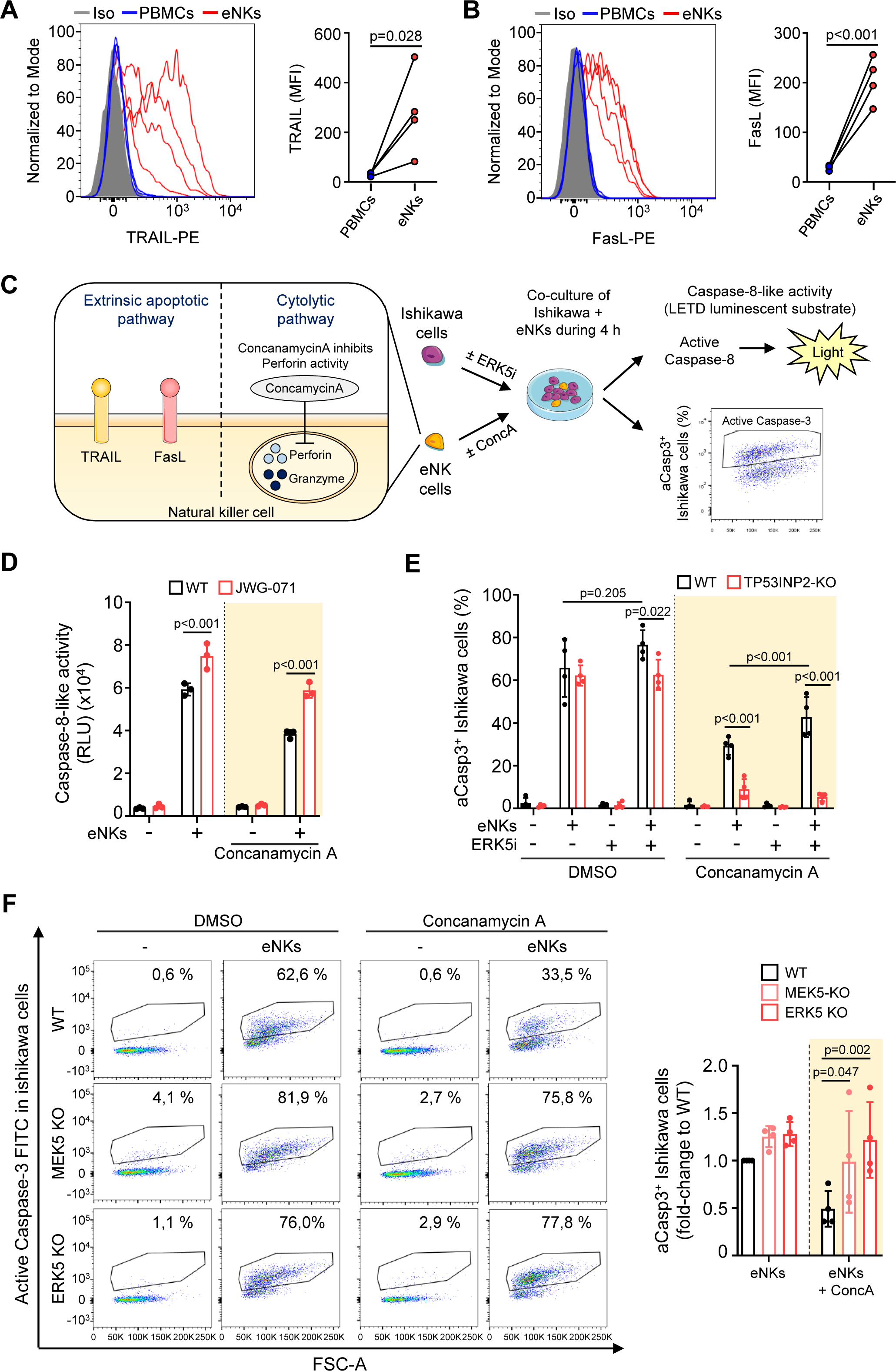
ERK5 inhibition or genetic deletion sensitizes Ishikawa EC cells to the cytotoxicity induced by the TRAIL and FasL expressed by eNK cells. **A-B**, Expanded NK cells express detectable TRAIL and FasL proteins at the membrane. **A**, TRAIL expression at the membrane of eNKs was detected by flow cytometry analysis. **B**, eNK cells were co-cultured with Ishikawa EC cells (4 h) and surface FasL was detected by flow cytometry. **C**, Schematic representation of the experiments performed co-culturing eNKs and EC cancer cells. eNKs from four different healthy donors were treated for 2 h with either vehicle (DMSO) or concanamycin A (yellow shading) to inhibit Perforin. Concanamycin A was maintained during the co-culture. **D**, ERK5i enhances caspase-8 activation induced by eNKs (white shading), and caspase-8 activation induced by the eNKs TRAIL/FasL pathway (yellow shading) in EC cells. **E**, TP53INP2 mediates EC cell sensitization to eNKs TRAIL/FasL pathway exerted by ERK5i. Active caspase-3 flow cytometry assay. Yellow shading histograms: TP53INP2-KO Ishikawa EC cells show impaired activation of caspase-3 in response to the eNK TRAIL/FasL cytotoxic pathway. Results are the mean ± SD of four independent biological samples (donors). **F**, MEK5 or ERK5 genetic deletion sensitize Ishikawa EC cells to the cytotoxicity induced by the eNK TRAIL/FasL cytotoxic pathway. Flow cytometry analysis for caspase-3 activation (cleaved caspase-3). Right histograms show the corresponding caspase-3 quantification. Yellow shading histograms show the results obtained for the eNK TRAIL/FasL cytotoxic pathway. Results are the mean ± SD of four independent biological samples (donors). **D, E and F**, Statistical significance was calculated using one-way ANOVA followed by Bonferroni multiple comparison test. Individual p-values are indicated in each panel.

Ishikawa EC cells were treated with ERK5 inhibitor or vehicle, exposed to human eNKs, and apoptosis was quantified by measuring either caspase-8 activity (luminescent substrate LETD) or cleaved caspase-3 (flow cytometry staining) (**Fig. 8C**). In EC cells, ERK5 inhibition enhanced eNK-induced caspase-8 activity even when perforin was inhibited with Concanamycin A (42) (**Fig. 8D**, yellow shading). Parallel experiments showed that ERK5i also tended to increase (but not significantly) eNKs-induced caspase-3 activity in Ishikawa cells, an effect which became particularly evident when the eNK cytolytic pathway was inhibited with Concanamycin A (**Fig. 8E**). These results suggest that ERK5 inhibition sensitizes EC cells to the extrinsic apoptosis induced by TRAIL and FasL expressed by NK cells. This effect was mediated by TP53INP2, since TP53INP2 genetic deletion mostly protected Ishikawa cells from apoptosis induced by eNK cell TRAIL/FasL (**Figure 8E**). Of note, the ERK5 inhibitor did not sensitize TP53INP2-KO cells to apoptosis induced by TRAIL/FasL expressed by eNKs (**Figure 8E**). Analogous experiments were carried out using MEK5-KO or ERK5-KO EC cells. Similar to ERK5 inhibition, genetic deletion of MEK5 or ERK5 increased eNK cell-induced apoptosis of EC cells, particularly when the cytolytic pathway was inhibited (**Fig. 8F**).

Together, our results point to the ERK5-TP53INP2 pathway as a negative regulator of NK-TRAIL/FasL-induced apoptosis in cancer cells, and ERK5 inhibition as a new strategy to improve the anticancer activity of NKs.

## Discussion

Extrinsic apoptosis is triggered by direct binding of DR ligands (TRAIL, TNFα and FasL) to their cognate receptors. In this study, we described a new signaling pathway involved in sensitization of cancer cells to the apoptosis induced by death receptor ligands. Hence, MAPK ERK5 phosphorylates and induces the ubiquitylation and proteasomal degradation of TP53INP2, a protein required for K63 ubiquitylation and full activation of caspase-8 in response to DR ligands.

Recent work has described an important role for TP53INP2 in regulating the extrinsic apoptotic pathway. TP53INP2 binds to caspase-8, acting as a scaffolding protein that facilitates the recruitment of the E-3 ligase TRAF6(5). TRAF6 mediates caspase-8 K63-linked ubiquitylation, required for caspase-8 oligomerization and full activation (30). Thus, ectopic overexpression of TP53INP2 sensitized cancer cells to DR ligands-induced apoptosis (5), suggesting modulation of TP53INP2 proteostasis as a new strategy to improve the apoptosis induced by DR ligands. Here, we showed that cellular TP53INP2 has a short half-life (4 hours), is degraded at the proteasome upon ubiquitylation, and that phosphorylation regulates its stability. Specifically, we found that ERK5 phosphorylates at least three residues (S49, S65 and T74) located at the N-terminal end of TP53INP2, between the LC3-interacting region (LIR) and the ubiquitin interacting motif (UIM) (5). Although we cannot discard the phosphorylation of other residues by ERK5, our results suggest that ERK5-mediated phosphorylation of S49, S65 and T74 is enough to induce TP53INP2 ubiquitylation and proteasomal degradation. Thus, it is likely that this multisite phosphorylation could allow the recruitment of the putative E3 ligase responsible for TP53INP2 ubiquitylation, yet to be discovered. In this regard, ERK5 is emerging as a protein kinase that regulates the stability of its substrates. For instance, ERK5 phosphorylates KLF2 transcription factor at three Ser/Thr-Pro residues (T171, S175 and S247), allowing the recruitment of FBXW7 substrate adaptor of the RING-finger E3 ubiquitin ligase Cullin-1, which ubiquitylates and induces proteasomal degradation of KLF2 (36). On the contrary, phosphorylation of c-MYC at Ser62 by ERK5 impairs the recruitment of the FBXW7 substrate adaptor, resulting in increased c-MYC protein stability (37). It will be important to investigate whether FBXW7 also regulates TP53INP2 ubiquitylation upon ERK5 phosphorylation.

We found that ERK5 pathway inhibition or genetic deletion sensitized cancer cells to DR ligands-induced apoptosis. This effect was mediated by the stabilization of TP53INP2 protein levels. Mechanistically, ERK5 inhibition impaired TP53INP2 proteasomal degradation, which resulted in elevated TP53INP2 protein levels and higher TP53INP2-Caspase 8 complex levels. In this scenario, elevated TP53INP2 levels could facilitate the ubiquitylation and full activation of caspase-8 in response to DRs activation. Whether TP53INP2 stability is also regulated by other oncogenic protein kinases, as the MEK5-ERK5 pathway does, deserves future investigation. If so, TP53INP2 proteostasis could emerge as a central node regulating sensitivity/resistance to DR agonist cytotoxicity, by integrating oncogenic signaling pathways involved in cancer cell survival. Of note, we cannot rule out other players contributing to the sensitization to DR ligands-induced apoptosis exerted by inhibition of the ERK5 pathway. In this regard, transcriptomic analysis identified minor but significant changes on FLIP (log_2_ FC -0,57) and XIAP (log_2_ FC -0,27) mRNA levels in response to ERK5 inhibition, two key proteins involved on resistance to DR ligands cytotoxicity (43). Future work is needed to investigate whether ERK5 inhibition impairs FLIP and XIAP protein level, as observed for the CDK1/2/9 and ERK5 multiple kinase inhibitor TG02 (44).

Among DR ligands, only TRAIL-receptor agonists (TRAs) have been developed as anticancer agents. Although FasL and TNFα show potent cytotoxicity in cancer cells, the use of these DR agonists in the clinic has been jeopardized by their severe toxicity in normal cells and tissues. Thus, TNF-α and FasL are cytotoxic towards hepatocytes, and induce lethal liver injuries in animal models (45,46). On the contrary, TRAIL selectively induces apoptosis in cancer cells (6). However, despite initial hopes, TRAs have shown limited therapeutic benefits in clinical trials due to inherent resistance of primary tumors to TRAIL-treatment (47). Consequently, much effort has been addressed to identify strategies to improve the anticancer activity of TRAs. Initially, strategies addressed specific resistance mechanisms driven by anti-apoptotic proteins such as IAPs and Bcl-2 family members, but they did not show benefit in clinical trials due to the heterogeneous nature of human cancers (48). Lately, it has been proposed the use of protein kinase inhibitors to remove resistance to TRAs-based therapies, given their ability to simultaneously block both oncogenic pathways and several proteins involved in this resistance (7). In this work, using cancer cell lines from different origin and known to be resistant to recombinant TRAIL, we demonstrated that pharmacologic inhibition of ERK5 is an effective strategy to improve TRAIL cytotoxicity, by stabilizing TP53IP2 protein levels. Interestingly, ERK5 inhibition or MEK5 genetic deletion also sensitized NSCLC KRAS mutated A549 cells to TRAIL cytotoxicity, a model where TRAIL induces cancer progression, invasion and metastasis instead of activating apoptosis (47,49). These preliminary results suggest that ERK5i switches the protumoral phenotype of TRAIL to an antitumoral (apoptotic) phenotype in NSCLC harboring activating KRAS mutations. We also found that ERK5i sensitized EC patient-derived xenografts (PDXOs, a highly relevant pre-clinical model) to rTRAIL toxicity. Since PDXOs maintain the genetic profile of EC subtypes predicting patient prognosis against treatments (32), our results support the use of ERK5 inhibitors to sensitize EC tumors to TRAs-based therapies. Future work is required to investigate whether this strategy might work for other cancer types.

Natural killer cells are critical players of the anti-tumor action of the immune system. Lately, NK cells have gained significant attention for cancer immunotherapy, since their cytotoxic activity does not require prior exposure to tumor antigens and they can efficiently kill tumor cells with minimal graft-versus-host disease (50). However, tumor cells frequently escape the immune attack due to inherent resistance mechanisms to apoptosis (51). Therefore, drugs that sensitize tumors to the NK cytotoxicity are gaining clinical relevance. Here, we provide evidence showing that ERK5 inhibition sensitizes EC cells to the extrinsic apoptosis induced by TRAIL/FasL expressed by eNKs. Our results agree with a previous report showing that ERK5 silencing sensitized leukemic cells to the cytotoxicity exerted by FasL-expressing NK cells (52), although the authors did not deep into the mechanistics of such effect. Our study points TP53INP2 protein as the key player in this process, since sensitization of EC cells to NK cells required TP53INP2 protein stabilization. Furthermore, the enhanced eNK-induced apoptosis induced by ERK5i is due to an effect on the TRAIL/FasL-induced extrinsic pathway rather than on intrinsic apoptosis induced by perforin and granzymes. However, ERK5 inhibitors might be relevant for improving the efficacy of NK-based immunotherapies, since NK cells switch from granzyme/perforin-mediated cancer cell death to TRAIL/FasL-induced cytotoxicity during subsequent tumor cell encounters (also called serial killing) (40). In this regard, it will be important to investigate whether modulation of the ERK5-TP53INP2 pathway also affects the expression of components of both cytolytic (granzymes/perforin) and TRAIL/FasL pathways in NK cells. Whether the results observed with our preparations of eNK cells might be extrapolated to different clinical adoptive immunotherapy protocols deserve attention (53).

The MEK5-ERK5 pathway regulates proliferation and survival of many solid cancers, including endometrial cancer (14), among others. MEK5 or ERK5 pharmacologic inhibitors have shown anticancer activity in xenograft models of several cancers, so ERK5 has been proposed as a new target to tackle several solid cancers. Of relevance for our work, ERK5 inhibitors sensitize tumor xenografts to standard chemotherapics such as 5-fluorouracil (54), doxorubicin (55), paclitaxel (14) or docetaxel (56), potentiating the intrinsic apoptosis exerted by chemotherapy treatments.

Here, we provide for the first time evidence supporting a role for ERK5 kinase activity in the extrinsic apoptotic pathway. Hence, we propose pharmacologic modulation of ERK5 kinase activity as an effective way to improve the anticancer efficacy of apoptosis activating drugs, and support further preclinical development of ERK5 inhibitors.

## Acknowledgments

S.E.G. and A.G.G: are recipients of a fellowship from FI-AGAUR (2020-FISDU-00575 and FI_B 00293, respectively). I.B.A. is recipient of a PIF fellowship from UAB. M.V.C is recipient of a PFU fellowship (Ref. FPU21/01999). B.V.M. is recipient of a PFIS fellowship (ref. FI21/00271). We thank Miguel Segura and Guillermo Yoldi for helpful discussions. We also thank the VHIR High Technology Unit (UAT) core facilities and staff, and the LP-CSIC/UAB Proteomic Laboratory for proteomics analyses. J.M.L. acknowledges support from Spanish Ministry of Economy and Competitiveness (MINECO, SAF2015-64237-R), Spanish Ministry of Science and Innovation (PID2019-107561RB-I00), and co-funded by the European Regional Development Fund (ERDF, “A way to make Europe/Investing in your future”). E.C. was founded by Fundación Científica Asociación Española Contra el Cáncer (AECC, GCTRA1804MATI) and Generalitat de Catalunya (2021-SGR-01157). A.Z. acknowledges support by research grants from MINECO (PID2019-106209RB-I00), CIBERDEM (Instituto de Salud Carlos III), and from an ICREA “Academia” Award (Generalitat de Catalunya). M.L.B. acknowledges support by research grant PID2019-110609RB-C21 from MCIN/AEI/ 10.13039/501100011033.

## Materials and Methods

### Reagents

ERK5 inhibitors JWG-071 (Merck), AX-15836 (MedChemExpress), and MEK5 inhibitors BIX02188 and BIX02189 (Selleckchem) were diluted in dimethyl sulfoxide (DMSO, Sigma). Pan-caspase inhibitor Q-VD(OMe)-OPh (APExBIO) was diluted in DMSO. Death-receptor agonists, TRAIL (Peprotech) and TNFα (Merck) were diluted in H_2_O 0.1% BSA, and anti-Fas activating antibody (Merck) was diluted in 50% glycerol in PBS.

### DNA constructs

The pEBG2T vectors encoding GST-tagged human ERK5 (WT) and ERK5 kinase-dead mutant (D200A) were a gift from Dr. P. Cohen (MRC Protein Phosphorylation Unit, Dundee, UK)(38). The pCMV5 vector encoding for HA-tagged constitutively active MEK5 (MEK5DD) was a gift from Dr. E. Nishida (Kyoto University, Japan) (57). The vectors encoding human N-Terminal FLAG-tagged TP53INP2 were generated by inserting the human TP53INP2 full-length coding sequence into a pCDNA3.1 vector, using the restriction sites NheI and XhoI. The FLAG-tagged human TP53INP2 mutants 3A (Ser49Ala, Ser65Ala, Thr74Ala) and 3K/R (Lys 165/187/204 mutated to Arg) were generated using the same strategy, inserting the synthetic sequences (ProteoGenix) into a pCDNA3.1 vector. The pCMV vector encoding 6xHis-tagged ubiquitin was a gift from Dr. A. C. Vertegaal (Leiden University, Netherlands) (58).

### Cells, cell culture and transfection

AN3CA, ARK1 and ARK2 endometrial cancer (EC) cells, HeLa cervical cancer cells, A549 non-small cell lung cancer cells, LnCaP prostate cancer cells and SK-N-AS neuroblastoma cells were purchased from ATCC. Ishikawa EC cells were from ECACC and were purchased from Sigma. HeLa, ARK1, ARK2 and A549 cells were maintained in Dulbecco’s Modified Eagle’s Medium (DMEM; Thermofisher) supplemented with 10% foetal bovine serum (FBS; Gibco) and 1% Penicillin/Streptomycin (Pen/Strep; Gibco). AN3CA were maintained in DMEM-F12 medium (ThermoFisher) supplemented with 10% FBS and 1% Pen/Strep. SK-N-AS cells were cultured in Iscove′s Modified Dulbecco′s medium (IMDM; Thermofisher) with 20% FBS (Gibco), 1% Pen/Strep (Gibco) and 1% Insulin-Transferrin-Selenium (Gibco). Ishikawa cells were maintained in Modified Eagle Medium (MEM; Thermofisher). LnCaP cells were maintained in Roswell Park Memorial Institute Medium (RPMI-1640; Gibco) supplemented with 10% FBS (Sigma), 1% Pen/Strep (Gibco), nonessential amino acids (NEAA; Sigma), sodium pyruvate (Gibco) and HEPES (Gibco). HEK293T cells were transfected using Polyethylenimine (PEI; Polysciences), and Ishikawa and AN3CA cells were transfected using Lipofectamine2000^TM^ (Thermofisher) as described before(59).

For 3D cultures of EC cells, a Matrigel (Corning) layer of approximately 1-2 mm in thickness was seeded per well in a 96-well plate. After solidification of the Matrigel layer at 37°C, 2500 Ishikawa cells were seeded per well in basal medium (DMEM/F-12, 1mM sodium pyruvate, 1% Pen/Strep) supplemented with 2% dextran-coated charcoal-stripped serum (Cytiva) and 3% Matrigel. 3D cultures were then left to grow during 5-7 days before performing the corresponding viability experiments.

### Generation of MEK5, ERK5 and TP53INP2 knockout cells by CRISPR/Cas9 technology

We used CRISPR/Cas9 technology to genetically delete the MEK5 gene (*MAP2K5*) in HeLa, Ishikawa and A549 cells, the ERK5 gene (MAPK7) in Ishikawa cells, and the TP53INP2 gene (*TP53INP2*) in Ishikawa cells. We used the following commercial kits from Santa Cruz technology: MEK5 (sc-401688-KO and sc-401688-HDR), ERK5 (sc-400891-KO and sc-400891-HDR) and TP53INP2 (sc-405261-KO-2 and sc-405261-HDR-2). Briefly, cells were co-transfected with the KO plasmid (containing the sgRNAs for the specific gene and the Cas9 endonuclease gene) and the HDR plasmid (which contains puromycin-resistance gene that is inserted into the cut sites of the target genes through a HDR mechanism). The KO cells acquiring the puromycin resistance gene were selected with 1 ug/ml of puromycin (Sigma) for ∼3 weeks. Single colonies were recovered, monitored for target gene expression, and further expanded and stored in liquid nitrogen.

### Cell lysis, immunoblotting and immunoprecipitation

For immunoblot analysis cells were lysed in ice-cold RIPA buffer supplemented with 50 mM NaF, 5 mM sodium-pyrophosphate and 1 mM sodium-orthovanadate. Cell lysates were sonicated and centrifuged at 12.000 g for 12 min at 4°C, and supernatants were stored at -20°C until use. Protein concentration was determined using Pierce™ Coomassie Plus reagent (Thermofisher). Proteins were resolved in SDS-PAGE gels and electrotransferred onto nitrocellulose membranes (Merck). After incubation with the appropriated primary antibody, detection was performed using horseradish peroxidase-conjugated secondary antibodies and enhanced chemiluminescence reagent (Bio-Rad). Primary and secondary antibodies used are given in **Suppl. Table 1**.

For immunoprecipitation, cells were lysed in ice-cold NP-40 buffer (50 mM Tris-HCl, 0.27M Sucrose, 1mM orthovanadate, 1mM EDTA, 1mM EGTA, 10 mM glycerol-phosphate disodium salt, 50 mM NaF, 5 mM sodium pyrophosphate, 0.5% (w/v) NP40, pH 7.5). Protein G-sepharose beads (Cytiva) bound to 2.5 ug of the corresponding antibody were incubated with 1 mg of cell lysate for 2h at 4°C with rotation. Then, immunoprecipitates were washed twice with NP-40 buffer supplemented with 150 mM NaCl, and once with kinase buffer (Tris-HCl 50 mM, 0.1 mM EGTA, pH 7.5). Proteins were then eluted with Laemmli buffer and boiled for 5 min at 95°C. Immunoprecipitation of cell lysates overexpressing FLAG-tagged or GST-tagged proteins were performed using anti-FLAG-M2 affinity beads (Merck) or glutathione-Sepharose beads (Cytiva).

### Cell viability and clonogenic assays

Cell viability was determined using the MTT (3-(4,5-dimethyl-2-thiazolyl)-2,5-diphenyl-2H-tetrazolium Blue, Sigma) reduction assay. For clonogenic assays, 1000 Ishikawa cells were seeded in 6-wells plates. Twenty-four hours later, cells were treated with the indicated treatments for 10 days and stained with 0.5% crystal violet solution. Colonies were counted using ImageJ (FiJi).

### Flow cytometry analysis of apoptosis

Apoptosis analysis was performed using an Annexin-V Apoptosis detection kit (Invitrogen). Briefly, treated cells were trypsinized and counted. 10^5^ cells were selected for the staining. Cells were washed with PBS, twice with the binding buffer, and finally incubated with Annexin-V-APC and PI for 15 minutes. Samples were then acquired on a BD LSRFortessa (BD Biosciencies) flow cytometer and analyzed with FlowJo software (vX.0.7, TreeStar).

### Caspase-8 activity assay

Cells were cultured in 96-well white plates with clear bottom (Corning). Twenty-four hours later, cells were treated, and caspase-8 activity was measured using the Caspase-Glo® 8 kit (Promega), which uses the specific caspase-8 substrate LETD sequence, following the manufacturer’s instructions. Luminescence was measured in a Spark microplate reader (TECAN).

### In vitro radiochemical ERK5 kinase activity assay

Lysates from HEK293 cells overexpressing FLAG-tagged wild typeTP53INP2 or the TP53INP2-3A were affinity-purified with anti-FLAG-M2 affinity beads (Merck). Then, beads containing the immunoprecipitated TP53INP2 proteins were incubated with 200 ng of active ERK5(38) in kinase buffer. Reactions (total volume 50 μL) were started by adding a mixture of 10 mM MgCl_2_, 200 μM [γ-^32^P]-ATP (400 cpm/pmol). Assays were carried out at 30°C for 15-30 min and terminated by adding 5x Laemmli buffer. Finally, samples were boiled for 5 min at 95°C, resolved by SDS/PAGE gel electrophoresis, and radioactivity detected by autoradiography.

### In vivo TP53INP2 ubiquitination assay

HEK293T cells overexpressing His-Tagged Ubiquitin, FLAG-TP53INP2 and/or GST-ERK5 and HA-MEK5DD (constitutively active), were resuspended in 1 ml of Buffer A (6M guanidinium-HCl, 100 mM Na2HPO4/NaH2PO4, 10 mM Imidazole, pH 8), sonicated for 1 minute, passed through a 25G syringe, and finally centrifuged for 15 min at 15.000 g. The resulting supernatants were incubated with 25 ul of Ni^2+^-NTA-agarose beads for 2 h at room temperature with rotation. Beads were successfully washed as follows: twice with Buffer A, four times with Buffer B (buffer A diluted 1:3 in buffer C (25 mM Tris-HCl, 20 mM Imidazole, pH 6.8), and twice with buffer C. Ubiquitylated proteins were eluted with Buffer C containing 250 mM Imidazole for 15 min at 37°C. After adding Laemmli buffer, eluted proteins were boiled for 5 min at 95°C, resolved in a SDS/PAGE gel and immunoblotted using anti-TP53INP2 antibody.

### Phosphoproteomic analysis

Purified FLAG-tagged TP53INP2 was incubated with 200 ng of active ERK5 in kinase buffer with 200 µM ATP for 30 min at 30°C, in a total reaction volume of 50 µl. The reaction was terminated by adding 5x Laemmli buffer, samples resolved in SDS/PAGE gel electrophoresis and stained with Coomassie Blue. The band corresponding to TP53INP2 was excised and digested with sequencing grade modified trypsin (Promega), using LFASP digestion protocol. Extracted peptides were analyzed by label-free liquid chromatography-tandem mass spectrometry (LC-MS/MS) at the LP-CSIC/UAB Service facility. LC-MS/MS was performed using an Agilent 1200 nanoflow system (Agilent Technologies) coupled to an LTQ-Orbitrap XL mass spectrometer (Thermo Fisher Scientific) equipped with a nanoelectrospray ion source (Proxeon Biosystems). Peptides were identified using Proteome Discoverer v1.4 and SwissProt human database, and analyzed using MASCOT database search (matrixscience.com) to identify phospho-Ser/Thr and phospho-Tyr.

### Generation of EC Patient-Derived Xenograft Organoids (PDXOs)

Each tumor tissue from EC PDXs was transferred into a 50-ml Falcon tube containing washing medium (DMEM/F12 supplemented with 1% L-glutamine, Penicillin-Streptomycin and 10 mM HEPES) and kept on ice until the start of the isolation. Tissues were minced using scalpels, followed by sedimentation with washing medium supplemented with 0.1% BSA during 2-3 min, to wash away any potential microorganisms or cell types not contributing to the establishment of organoids. Enzymatic digestion was done with 1.25 U/ml dispase II (Merck), 0.4 mg/ml collagenase from Clostridium histolyticum type V (Merck), and ROCK inhibitor Y-27632, on an orbital shaker for 60 min at 37 °C. Undigested tissue was discarded using a 100 μm cell strainer, and red blood cell lysis buffer (Merck) was subsequently added to the pellet. Single cells were then resuspended in BME (basement membrane extract, Bio-Techne) and seeded (20 μl droplets) onto non-treated 12-well culture plates. The suspension of organoids and BME was solidified at 37 °C for 10-15 min before covering each well with 750 μl of standard organoid medium containing washing medium supplemented with 100 ng/ml recombinant human EGF (Bio-Techne), recombinant human Noggin (100 ng/ml, Stemcell Technologies), B-27 supplement (ThermoFisher), 10 μM Y-27632 (Miltenyi Biotec), 100 nM SB202190 (Merck), 10 nM 17-β Estradiol (Merck), 10 μM Nicotinamide (Merck), 500 nM A83-01 (Merck) and N-acetyl-L-cysteine 1.25mM (Merck).

### Viability assays in 3D cultures and EC PDXOs

Cells were seeded in 96-well white plates with clear bottom (Corning). When the 3D cultures or organoids were formed, they were treated for the indicated times and viability was assessed using the CellTiter-Glo® 3D Cell viability kit (Promega), following the manufacturer’s instructions. The CellTiter-Glo® 3D Cell viability kit measures the intracellular ATP content of the 3D cultures which is directly related to the viability of the cells. Luminescence was measured in a Spark microplate reader (TECAN). In parallel experiments, viability measurement was performed using a LIVE/DEAD viability/cytotoxicity kit assay (Thermofisher), as described before (59).

### EC PDXOs immunofluorescence microscopy

Organoids were extracted from their BME matrix by adding ice-cold cell recovery solution (Corning) for 30-60 minutes. This procedure enables the dissolution of BME without damaging the organoids, and allows an optimal penetration of the antibody. Then, organoids were fixed with 4% (w/v) paraformaldehyde at 4°C for 45 min. Throughout the protocol, samples were washed extensively to avoid background or loss of signal. Primary antibodies E-cadherin (Abcam), Vimentin (Abcam) and EpCAM (Cell Signalling) were incubated overnight at 4°C on a horizontal shaker (40 rpm). Secondary antibodies Mouse IgG Alexa Fluor 555 (Abcam), Rabbit IgG Alexa Fluor 488 (Abcam) and the Phalloidin-iFluor 647 (Abcam) were added to each well and incubated overnight in the same conditions. Antibodies used in this process are listed in **Suppl. Table 2**. Optical clearing was accomplished by resuspending the organoids with homemade fructose-glycerol clearing solution. For slide preparation, a slice of double-sided tape was added on both sides of the slide to lift the coverslip, thereby maintaining the 3D structure of the organoids. The slide was then visualized by fluorescence confocal microscopy (Zeiss LSM980).

### RNA-seq analysis

Ishikawa cells were treated with either vehicle or JWG-071 for 24 h, and then lysed for RNA purification. RNA was purified from Ishikawa cells using RNeasy kit (Qiagen). RNA quality was evaluated using a Bioanalyzer 2100 (Agilent), and samples with a RIN value higher than 7 were selected. Then, libraries were prepared using TruSeq Kit (Illumina). RNA-sequencing (30 million reads per sample) was carried out at the DNBseq™ NGS technology platform with the Ilumina HiSeq4000 at the Beijing Genomics Institute (BGI). Finally, functional enrichment analyses were performed using Dr. Tom software (BGI), the gene set enrichment analysis (GSEA) software and the gene ontology (GO) collection (MSigDB v7.1).

### Natural killer cell amplification and immunophenotyping characterization

Freshly isolated PBMCs (peripheral blood mononuclear cells) from healthy human donors were co-cultured with irradiated (40 Gy) RPMI 8866 B lymphoblastoid cell line in a ratio 3:1 (3×10^6^ PBMCs and 1×10^6^ 8866 cells per well in a 12 well-plate) in RPMI 1640 GlutaMax (Thermo Fisher Scientific) supplemented with penicillin (100 U/ml), streptomycin (100 μg/ml), sodium pyruvate (1 mM), and 10% FBS. Cultures were split and fed every 3-4 days, adding IL-2 (200 U/ml) from day 7. On day 12, cultures were harvested and expanded natural killer (eNK) cells (CD56^+^/CD3^-^) were purified (Human NK Cell Isolation kit, Miltenyi). Expression of cytotoxic markers (Granzyme B, Perforin, TRAIL and FasL), activating (NKG2D, NKp46, NKp30, CD16) and inhibitory (KIRs, ILT2, NKG2A) NK cell receptors (NKR), as well as the CD57 marker was assessed. Briefly, PBMC or eNK cells were treated with blocking buffer (2% FBS, 2mM EDTA, 10 µg/mL aggregated human IgG in PBS) and stained with the indicated labelled antibodies listed in Supplementary table 2. For intracellular staining, cells were fixed and permeabilized with Fix/Perm kit (BD Biosciences) and stained for intracellular antigens. Samples were acquired on a BD-LSRII (BD Biosciences) and analyzed with FlowJo software (vX.0.7, TreeStar). Antibodies used in this process are listed in **Suppl. Table 3**.

### Natural killer cell cytotoxicity assay

Ishikawa cells were pre-treated with either JWG-071 or vehicle (0.05% v/v DMSO) for 12h and detached. In parallel, eNK cells were pre-treated with 1 µM Concanamycin-A (MedChemExpress) or vehicle (0.1% v/v DMSO) for 2 hours. eNK cells were co-cultured with Ishikawa EC cells at 1:5 (effector:target) ratio in the presence of 1 µM Concanamycin A or vehicle for 4 hours in U-bottom 96-well plates (Corning), and active-caspase-3 in Ishikawa cells was analyzed by flow cytometry. Briefly, samples were stained with mAbs specific for CD45-BV510 (clone HI30) and EpCAM-AF647 (clone 9C4), together with Near IR-fluorescent reactive dye (Invitrogen) to exclude dead cells, permeabilized with wash/perm buffer, stained with anti-cleaved caspase-3-FITC antibody (BD Pharmingen, clone C92-605), acquired on a BD-LSRII flow cytometer (BD Biosciences) and analyzed with FlowJo software (vX.0.7, TreeStar). Percentages of active Caspase 3+ cells were assessed in viable Ishikawa cells (EpCAM^+^/CD45^-^). Antibodies used in this process are listed in **Suppl. Table 3**.

### Schematics and statistical analysis

The schemes in figures 6F, 8C and 9 were created with PowerPoint (Microsoft). Individual data points are displayed in all the bar plots. Data are presented as the mean ± standard deviation (S.D.) of at least n=3 independent experiments. The statistical tests used are reported in the figure legends. Significance was established using one-way ANOVA followed by Bonferroni multiple comparison test, two-tailed Student’s test or two-way ANOVA followed by Tukey multiple comparison test (individual p-values are represented in each figure). All the analyses were performed with GraphPad Prism 8.

### Data availability

The RNA sequencing datasets generated in this study have been deposited in the GEO database repository under accession code (pending). The remaining data are available within the Article, Supplementary Information or Dataset files.

## References

1. Zou H, Li Y, Liu X, Wang X. An APAF-1.cytochrome c multimeric complex is a functional apoptosome that activates procaspase-9. J Biol Chem. 1999;274(17):11549–56.

2. Medema JP, Scaffidi C, Krammer PH, Peter ME. Bcl-xL acts downstream of caspase-8 activation by the CD95 death-inducing signaling complex. J Biol Chem. 1998;273(6):3388–93.

3. Jin Z, Li Y, Pitti R, Lawrence D, Pham VC, Lill JR, et al. Cullin3-based polyubiquitination and p62-dependent aggregation of caspase-8 mediate extrinsic apoptosis signaling. Cell. 2009;137(4):721–35.

4. He L, Wu X, Siegel R, Lipsky PE. TRAF6 regulates cell fate decisions by inducing caspase 8-dependent apoptosis and the activation of NF-kappaB. J Biol Chem. 2006;281(16):11235–49.

5. Ivanova S, Polajnar M, Narbona-Perez AJ, Hernandez-Alvarez MI, Frager P, Slobodnyuk K, et al. Regulation of death receptor signaling by the autophagy protein TP53INP2. EMBO J. 2019;38(10).

6. Ashkenazi A, Dixit VM. Apoptosis control by death and decoy receptors. Curr Opin Cell Biol. 1999;11(2):255–60.

7. Montinaro A, Walczak H. Harnessing TRAIL-induced cell death for cancer therapy: a long walk with thrilling discoveries. Cell Death Differ. 2022.

8. Drew BA, Burow ME, Beckman BS. MEK5/ERK5 pathway: the first fifteen years. Biochim Biophys Acta. 2012;1825(1):37–48.

9. Kato Y, Kravchenko VV, Tapping RI, Han J, Ulevitch RJ, Lee JD. BMK1/ERK5 regulates serum-induced early gene expression through transcription factor MEF2C. EMBO J. 1997;16(23):7054–66.

10. Gomez N, Erazo T, Lizcano JM. ERK5 and Cell Proliferation: Nuclear Localization Is What Matters. Front Cell Dev Biol. 2016;4:105.

11. Stecca B, Rovida E. Impact of ERK5 on the Hallmarks of Cancer. Int J Mol Sci. 2019;20(6).

12. Rovida E, Di Maira G, Tusa I, Cannito S, Paternostro C, Navari N, et al. The mitogen-activated protein kinase ERK5 regulates the development and growth of hepatocellular carcinoma. Gut. 2015;64(9):1454–65.

13. Montero JC, Ocana A, Abad M, Ortiz-Ruiz MJ, Pandiella A, Esparis-Ogando A. Expression of Erk5 in early stage breast cancer and association with disease free survival identifies this kinase as a potential therapeutic target. PLoS One. 2009;4(5):e5565.

14. Dieguez-Martinez N, Espinosa-Gil S, Yoldi G, Megias-Roda E, Bolinaga-Ayala I, Vinas-Casas M, et al. The ERK5/NF-kappaB signaling pathway targets endometrial cancer proliferation and survival. Cell Mol Life Sci. 2022;79(10):524.

15. McCracken SR, Ramsay A, Heer R, Mathers ME, Jenkins BL, Edwards J, et al. Aberrant expression of extracellular signal-regulated kinase 5 in human prostate cancer. Oncogene. 2008;27(21):2978–88.

16. Pereira DM, Rodrigues CMP. Targeted Avenues for Cancer Treatment: The MEK5-ERK5 Signaling Pathway. Trends Mol Med. 2020;26(4):394–407.

17. Pi X, Yan C, Berk BC. Big mitogen-activated protein kinase (BMK1)/ERK5 protects endothelial cells from apoptosis. Circ Res. 2004;94(3):362–9.

18. Girio A, Montero JC, Pandiella A, Chatterjee S. Erk5 is activated and acts as a survival factor in mitosis. Cell Signal. 2007;19(9):1964–72.

19. Aza-Blanc P, Cooper CL, Wagner K, Batalov S, Deveraux QL, Cooke MP. Identification of modulators of TRAIL-induced apoptosis via RNAi-based phenotypic screening. Mol Cell. 2003;12(3):627–37.

20. Borges J, Pandiella A, Esparis-Ogando A. Erk5 nuclear location is independent on dual phosphorylation, and favours resistance to TRAIL-induced apoptosis. Cell Signal. 2007;19(7):1473–87.

21. Lane D, Goncharenko-Khaider N, Rancourt C, Piche A. Ovarian cancer ascites protects from TRAIL-induced cell death through alphavbeta5 integrin-mediated focal adhesion kinase and Akt activation. Oncogene. 2010;29(24):3519–31.

22. Pallares J, Martinez-Guitarte JL, Dolcet X, Llobet D, Rue M, Palacios J, et al. Abnormalities in the NF-kappaB family and related proteins in endometrial carcinoma. J Pathol. 2004;204(5):569–77.

23. Wang J, Erazo T, Ferguson FM, Buckley DL, Gomez N, Munoz-Guardiola P, et al. Structural and Atropisomeric Factors Governing the Selectivity of Pyrimido-benzodiazipinones as Inhibitors of Kinases and Bromodomains. ACS ChemBiol. 2018;13(9):2438–48.

24. Lin EC, Amantea CM, Nomanbhoy TK, Weissig H, Ishiyama J, Hu Y, et al. ERK5 kinase activity is dispensable for cellular immune response and proliferation. Proc Natl Acad Sci U S A. 2016;113(42):11865–70.

25. Tatake RJ, O’Neill MM, Kennedy CA, Wayne AL, Jakes S, Wu D, et al. Identification of pharmacological inhibitors of the MEK5/ERK5 pathway. Biochem Biophys Res Commun. 2008;377(1):120–5.

26. Gatsinzi T, Ivanova EV, Iverfeldt K. TRAIL resistance in human neuroblastoma SK-N-AS cells is dependent on protein kinase C and involves inhibition of caspase-3 proteolytic processing. J Neurooncol. 2012;109(3):503–12.

27. Poondla N, Chandrasekaran AP, Heese K, Kim KS, Ramakrishna S. CRISPR-mediated upregulation of DR5 and downregulation of cFLIP synergistically sensitize HeLa cells to TRAIL-mediated apoptosis. Biochem Biophys Res Commun. 2019;512(1):60–5.

28. Reis CR, Chen PH, Bendris N, Schmid SL. TRAIL-death receptor endocytosis and apoptosis are selectively regulated by dynamin-1 activation. Proc Natl Acad Sci U S A. 2017;114(3):504–9.

29. Chen X, Thakkar H, Tyan F, Gim S, Robinson H, Lee C, et al. Constitutively active Akt is an important regulator of TRAIL sensitivity in prostate cancer. Oncogene. 2001;20(42):6073–83.

30. Tummers B, Green DR. Caspase-8: regulating life and death. Immunol Rev. 2017;277(1):76–89.

31. Wang X, Merritt AJ, Seyfried J, Guo C, Papadakis ES, Finegan KG, et al. Targeted deletion of mek5 causes early embryonic death and defects in the extracellular signal-regulated kinase 5/myocyte enhancer factor 2 cell survival pathway. MolCell Biol. 2005;25(1):336–45.

32. Berg HF, Hjelmeland ME, Lien H, Espedal H, Fonnes T, Srivastava A, et al. Patient-derived organoids reflect the genetic profile of endometrial tumors and predict patient prognosis. Commun Med (Lond). 2021;1:20.

33. Villafranca-Magdalena B, Masferrer-Ferragutcasas C, Lopez-Gil C, Coll-de la Rubia E, Rebull M, Parra G, et al. Genomic Validation of Endometrial Cancer Patient-Derived Xenograft Models as a Preclinical Tool. Int J Mol Sci. 2022;23(11).

34. Mauvezin C, Orpinell M, Francis VA, Mansilla F, Duran J, Ribas V, et al. The nuclear cofactor DOR regulates autophagy in mammalian and Drosophila cells. EMBO Rep. 2010;11(1):37–44.

35. Sala D, Ivanova S, Plana N, Ribas V, Duran J, Bach D, et al. Autophagy-regulating TP53INP2 mediates muscle wasting and is repressed in diabetes. J Clin Invest. 2014;124(5):1914–27.

36. Brown HA, Williams CAC, Zhou H, Rios-Szwed D, Fernandez-Alonso R, Mansoor S, et al. An ERK5-KLF2 signalling module regulates early embryonic gene expression and telomere rejuvenation in stem cells. Biochem J. 2021;478(23):4119–36.

37. Vaseva AV, Blake DR, Gilbert TSK, Ng S, Hostetter G, Azam SH, et al. KRAS Suppression-Induced Degradation of MYC Is Antagonized by a MEK5-ERK5 Compensatory Mechanism. Cancer Cell. 2018;34(5):807–22 e7.

38. Erazo T, Moreno A, Ruiz-Babot G, Rodriguez-Asiain A, Morrice NA, Espadamala J, et al. Canonical and kinase activity-independent mechanisms for extracellular signal-regulated kinase 5 (ERK5) nuclear translocation require dissociation of Hsp90 from the ERK5-Cdc37 complex. Mol Cell Biol. 2013;33(8):1671–86.

39. Mody N, Campbell DG, Morrice N, Peggie M, Cohen P. An analysis of the phosphorylation and activation of extracellular-signal-regulated protein kinase 5 (ERK5) by mitogen-activated protein kinase kinase 5 (MKK5) in vitro. Biochem J. 2003;372(Pt 2):567–75.

40. Prager I, Watzl C. Mechanisms of natural killer cell-mediated cellular cytotoxicity. J Leukoc Biol. 2019;105(6):1319–29.

41. Duan S, Guo W, Xu Z, He Y, Liang C, Mo Y, et al. Natural killer group 2D receptor and its ligands in cancer immune escape. Mol Cancer. 2019;18(1):29.

42. Kataoka T, Shinohara N, Takayama H, Takaku K, Kondo S, Yonehara S, et al. Concanamycin A, a powerful tool for characterization and estimation of contribution of perforin- and Fas-based lytic pathways in cell-mediated cytotoxicity. J Immunol. 1996;156(10):3678–86.

43. Lemke J, von Karstedt S, Zinngrebe J, Walczak H. Getting TRAIL back on track for cancer therapy. Cell Death Differ. 2014;21(9):1350–64.

44. Alvarez-Fernandez S, Ortiz-Ruiz MJ, Parrott T, Zaknoen S, Ocio EM, San Miguel J, et al. Potent antimyeloma activity of a novel ERK5/CDK inhibitor. Clin Cancer Res. 2013;19(10):2677–87.

45. Ogasawara J, Watanabe-Fukunaga R, Adachi M, Matsuzawa A, Kasugai T, Kitamura Y, et al. Lethal effect of the anti-Fas antibody in mice. Nature. 1993;364(6440):806–9.

46. Costelli P, Aoki P, Zingaro B, Carbo N, Reffo P, Lopez-Soriano FJ, et al. Mice lacking TNFalpha receptors 1 and 2 are resistant to death and fulminant liver injury induced by agonistic anti-Fas antibody. Cell Death Differ. 2003;10(9):997–1004.

47. von Karstedt S, Montinaro A, Walczak H. Exploring the TRAILs less travelled: TRAIL in cancer biology and therapy. Nat Rev Cancer. 2017;17(6):352–66.

48. Cristofanon S, Fulda S. ABT-737 promotes tBid mitochondrial accumulation to enhance TRAIL-induced apoptosis in glioblastoma cells. Cell Death Dis. 2012;3(11):e432.

49. Pal S, Amin PJ, Sainis KB, Shankar BS. Potential Role of TRAIL in Metastasis of Mutant KRAS Expressing Lung Adenocarcinoma. Cancer Microenviron. 2016;9(2-3):77–84.

50. Liu S, Galat V, Galat Y, Lee YKA, Wainwright D, Wu J. NK cell-based cancer immunotherapy: from basic biology to clinical development. J Hematol Oncol. 2021;14(1):7.

51. Sordo-Bahamonde C, Lorenzo-Herrero S, Payer AR, Gonzalez S, Lopez-Soto A. Mechanisms of Apoptosis Resistance to NK Cell-Mediated Cytotoxicity in Cancer. Int J Mol Sci. 2020;21(10).

52. Charni S, Aguilo JI, Garaude J, de Bettignies G, Jacquet C, Hipskind RA, et al. ERK5 knockdown generates mouse leukemia cells with low MHC class I levels that activate NK cells and block tumorigenesis. J Immunol. 2009;182(6):3398–405.

53. Lopez-Botet M, De Maria A, Muntasell A, Della Chiesa M, Vilches C. Adaptive NK cell response to human cytomegalovirus: Facts and open issues. Semin Immunol. 2023;65:101706.

54. Pereira DM, Simoes AE, Gomes SE, Castro RE, Carvalho T, Rodrigues CM, et al. MEK5/ERK5 signaling inhibition increases colon cancer cell sensitivity to 5-fluorouracil through a p53-dependent mechanism. Oncotarget. 2016;7(23):34322–40.

55. Shukla A, Miller JM, Cason C, Sayan M, MacPherson MB, Beuschel SL, et al. Extracellular signal-regulated kinase 5: a potential therapeutic target for malignant mesotheliomas. ClinCancer Res. 2013;19(8):2071–83.

56. Yang Q, Liao L, Deng X, Chen R, Gray NS, Yates JR, III, et al. BMK1 is involved in the regulation of p53 through disrupting the PML-MDM2 interaction. Oncogene. 2012.

57. Kondoh K, Terasawa K, Morimoto H, Nishida E. Regulation of nuclear translocation of extracellular signal-regulated kinase 5 by active nuclear import and export mechanisms. MolCell Biol. 2006;26(5):1679–90.

58. Schimmel J, Larsen KM, Matic I, van Hagen M, Cox J, Mann M, et al. The ubiquitin-proteasome system is a key component of the SUMO-2/3 cycle. Mol Cell Proteomics. 2008;7(11):2107–22.

59. Gamez-Garcia A, Bolinaga-Ayala I, Yoldi G, Espinosa-Gil S, Dieguez-Martinez N, Megias-Roda E, et al. ERK5 Inhibition Induces Autophagy-Mediated Cancer Cell Death by Activating ER Stress. Front Cell Dev Biol. 2021;9:742049.

