## Supplementary material for "MAP kinase ERK5 modulates cancer cell sensitivity to extrinsic apoptosis induced by death-receptor agonists and Natural Killer cells": Supplentary Figures Tables and data

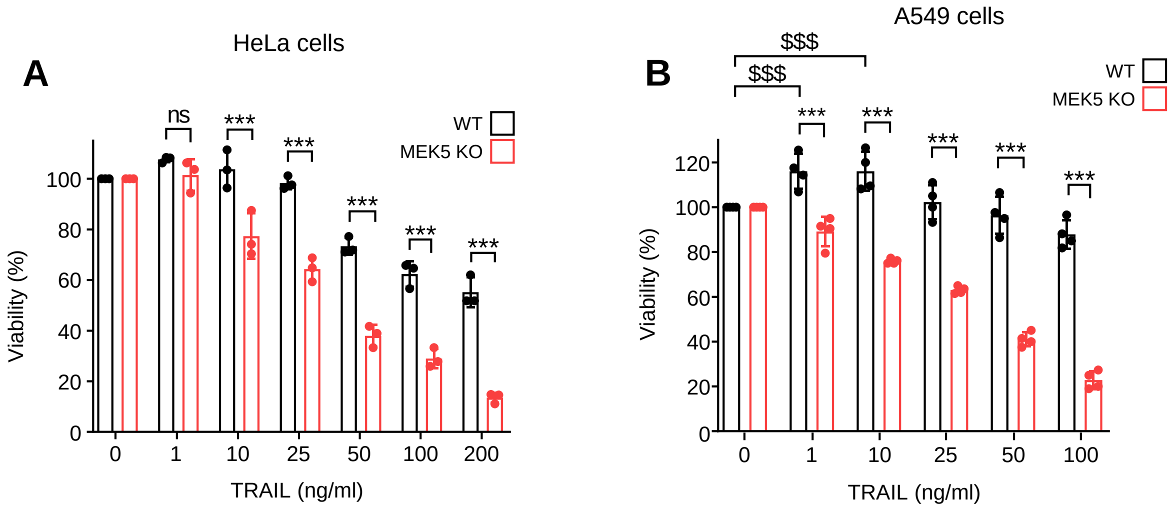


**Supplementary Figure 1. MEK5 genetic deletion confers increased sensitivity to TRAIL-induced toxicity in cervical cancer Hela and NSCLC A549 cells. A.** Cervical carcinoma (HeLa) or **B.** NSCLC cells (A549) were treated with TRAIL for 24 h and cell viability was analyzed. ***p < 0.001 (one-way ANOVA followed by Bonferroni multiple comparison test).

| Patient | Age | Histology | Grade | FIGO | p53 | MSH6 | PMS2 | POLE | TCGA |
|---|---|---|---|---|---|---|---|---|---|
| 440 | 75 | Endometrioid | 2 | II | WT | WT | Mut | WT | MSI |
| 1297 | 69 | Endometrioid | 3 | Ib | Mut | WT | Mut | Mut | POLE |

**Supplementary Figure 2. Clinicopathological features of endometrial cancer patients used for PDX-Os generation.** WT: wild-type; MSH6: MutS Homolog 6; PMS2: PMS1 Homolog 2; POLE: polymerase epsilon; MSI: microsatellite instability.

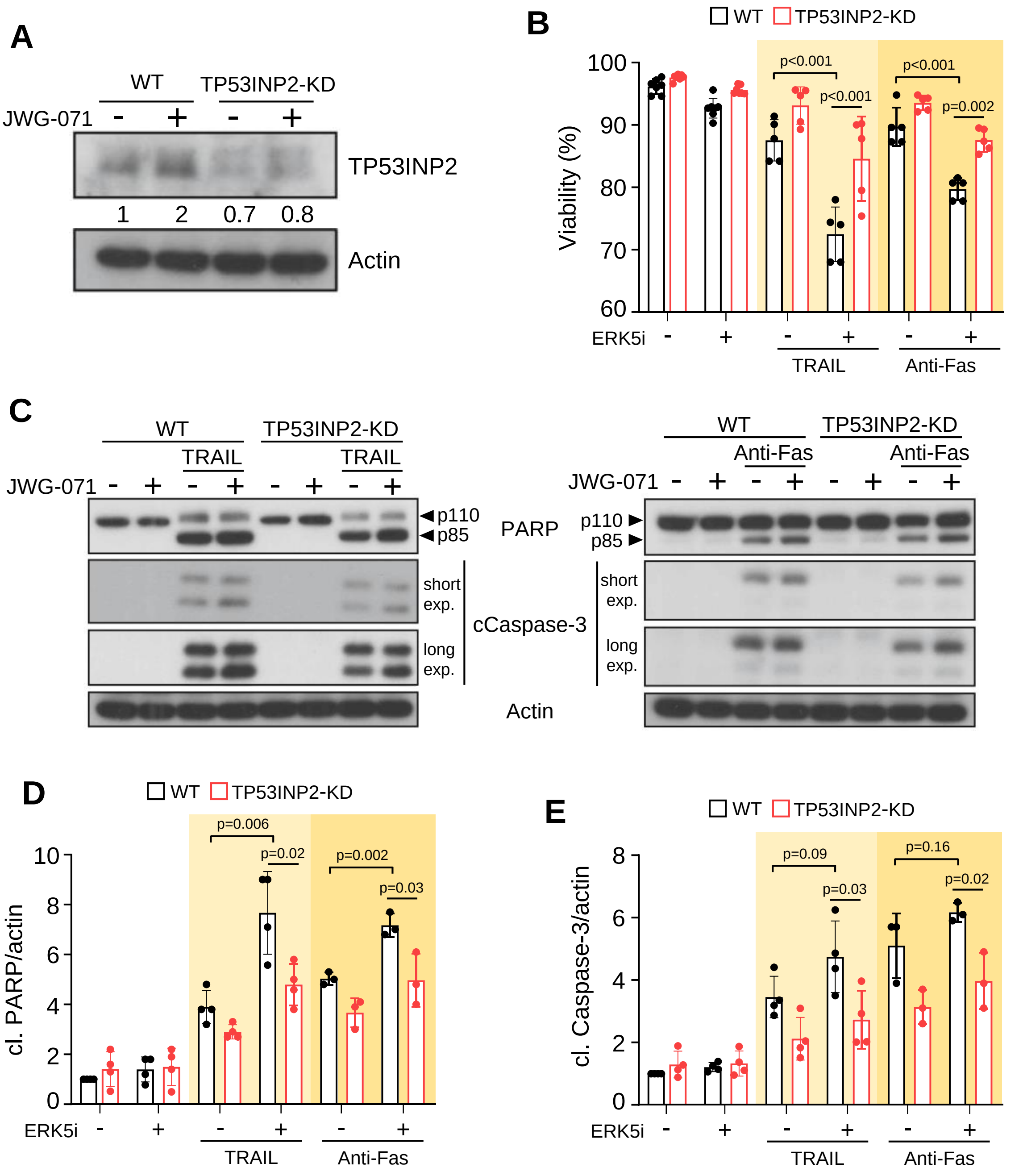


**Supplementary Figure 3. ERK5 inhibition sensitizes cervical cancer cells to DR agonists by increasing TP53INP2 protein levels. A**, ERK5 inhibition upregulates TP53INP2 protein levels in cervical cancer cells. HeLa cells were treated for 24 h with either vehicle or 5 µM JWG-071. TP53INP2 protein levels were determined by immunoblot analysis. **B-E,** TP53INP2 mediates the sensitization to DR agonists exerted by the ERK5i JWG-071 in HeLa cells. **B**, HeLa wild type or TP53INP2-KD cells (were TP53INP2 was knocked-down) were pre-treated 5 µM JWG-071 (16 h), treated with 50 ng/ml TRAIL or 100 ng/ml Anti-Fas activating antibody for 24 h, and cell viability was assessed by annexin V and PI staining. **C**, Activation of apoptosis was measured by immunoblot analysis of cleaved caspase-3 and PARP. Cells were pre-treated with JWG-071 (12 h) and further treated with 50 ng/ml TRAIL or 100 ng/ml Anti-Fas activating antibody for 4 h. **D-E,** Quantification of protein levels of cleaved PARP (**D**) and cleaved caspase-3 (**E**) after TRAIL and anti-Fas treatment. Statistical significance in B, D and E was calculated using two-tailed unpaired t-test. Individual p-values are indicated in each panel.

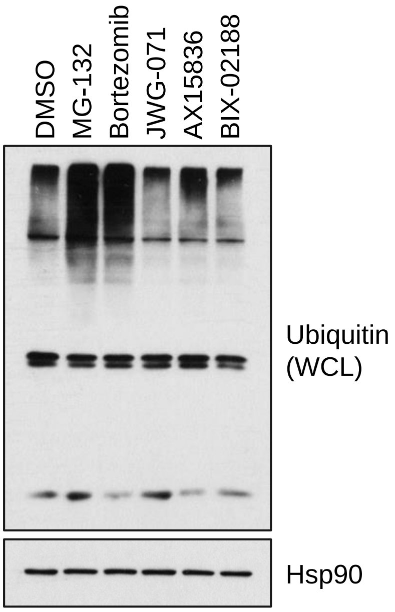


**Supplementary Figure 4. MEK5-ERK5 pathway inhibitors do not inhibit the proteasome.** Ishikawa cells were treated with proteasome inhibitors (MG-132 or Bortezomib) or ERK5 (JWG-071 and AX15836) and MEK5 (BIX02188) inhibitors for 8 h. Then, cells were lysed and global protein ubiquitylation was determined by immunoblotting for Ubiquitin.

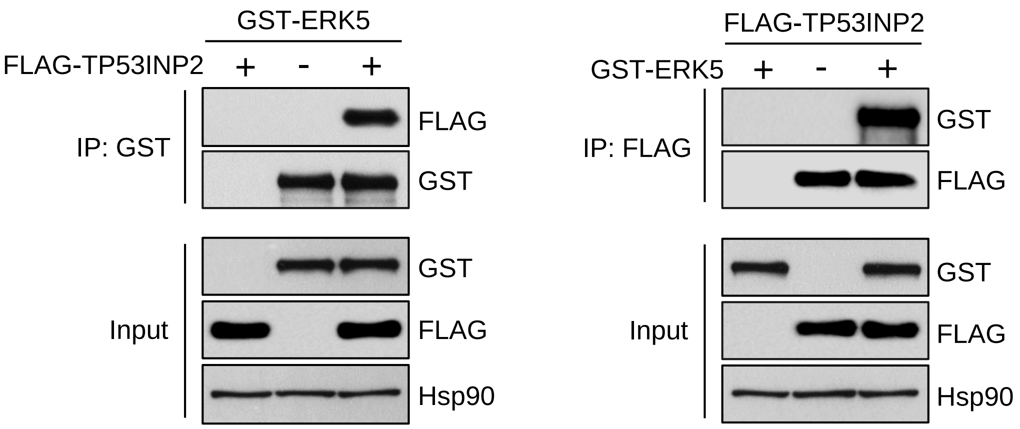


**Supplementary Figure 5. ERK5 interacts with TP53INP2.** HEK293T cells were transfected with a vector encoding for FLAG-tagged TP53INP2 alone or together with GST-tagged ERK5. Forty-eight hours later, cells were lysed in NP40 buffer and FLAG-TP53INP2 or GST-ERK5 were affinity-purified using ANTI-FLAG® M2 Affinity Beads or glutathione-sepharose, respectively. Immune complexes were immunoblotted for ERK5 and TP53INP2.

**A**

| Sequence | #PSMs | Modifications | MH+ [Da] | ERK5 | Empty |
|---|---|---|---|---|---|
| SKNQS**S**FIYQPCQR | 10 | S6(Phospho); C12(Carbamidomethyl) | 1822,79 | x | x |
| LSSLFFSTP**S**PPEDPDCPR | 20 | S10(Phospho); C17(Carbamidomethyl) | 2228,96 | x | x |
| R**S**KNQSSFIYQPCQR | 1 | S2(Phospho); C13(Carbamidomethyl) | 1978,90 | | x |
| NQS**S**FIYQPCQR | 3 | S4(Phospho); C10(Carbamidomethyl) | 1607,67 | x | x |
| LSSLFFS**T**P**S**PPEDPDCPR | 7 | T8(Phospho); S10(Phospho); C17(Carbamidomethyl) | 2308,93 | x | x |

**B**

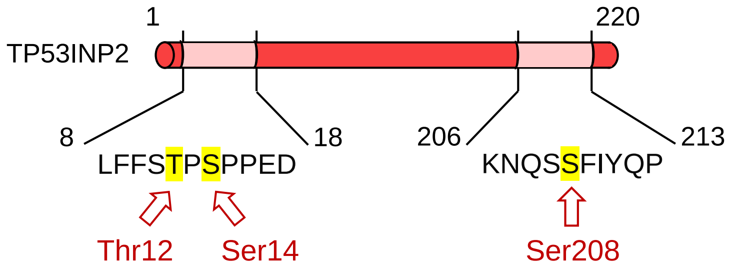


**Supplementary Figure 6. TP53INP2 protein is constitutively phosphorylated in Thr14, Ser14 and Ser208 in cells. A.** Protein TP53INP2 (Q81HX6) phosphorylated peptides identified by LC-MS/MS, and the graphical representation of its corresponding residues in the original TP53INP2 protein sequence (**B**).

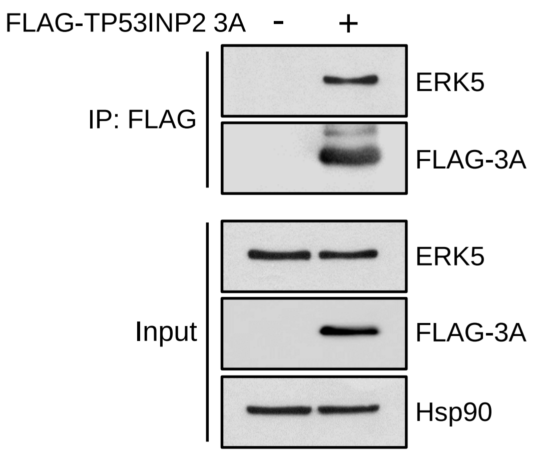


**Supplementary Figure 7. The TP53INP2 phospho-deficient mutant retains the ability to interact with ERK5.** HEK293T cells were transfected with a vector encoding for FLAG-tagged TP53INP2-3A. Forty-eight hours later, cells were lysed in NP40 buffer and FLAG-TP53INP2 was affinity-purified using ANTI-FLAG® M2 Affinity Beads. Immune complexes were immunoblotted for endogenous ERK5 and TP53INP2.

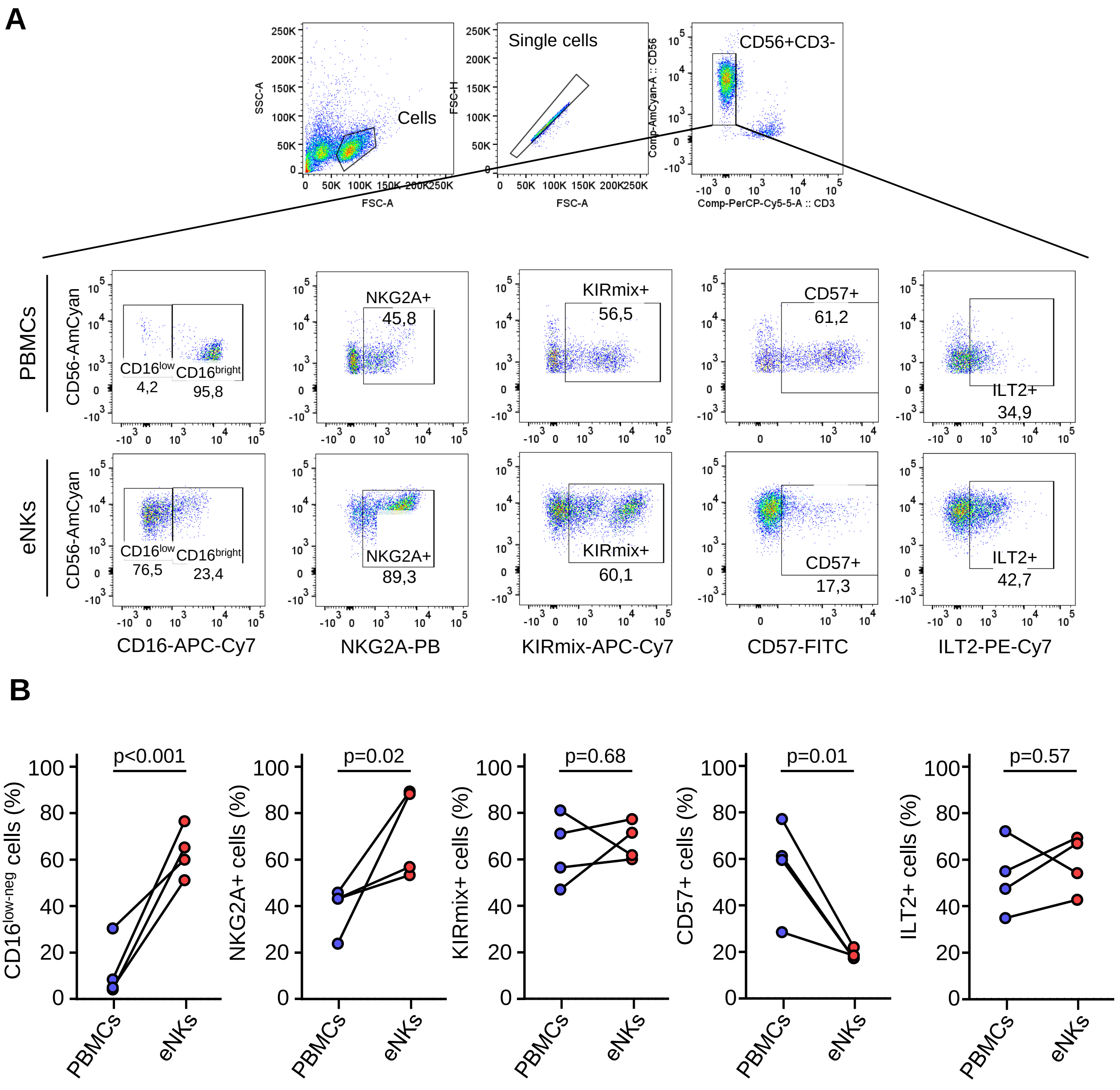


**Supplementary Figure 8. Comparative phenotypic analysis of fresh and eNK cells. A-B.** eNK cells and PBMCs from four healthy donors were stained for different NK cell surface markers (NKG2A, KIR, ILT2, CD16, CD57), analyzed and gated on forward and side scatter (FSC/SSC) and CD56+ /CD3- cells. Representative flow cytometry dot plots **(A)** and graphs of the results from samples of four donors **(B)** are depicted.

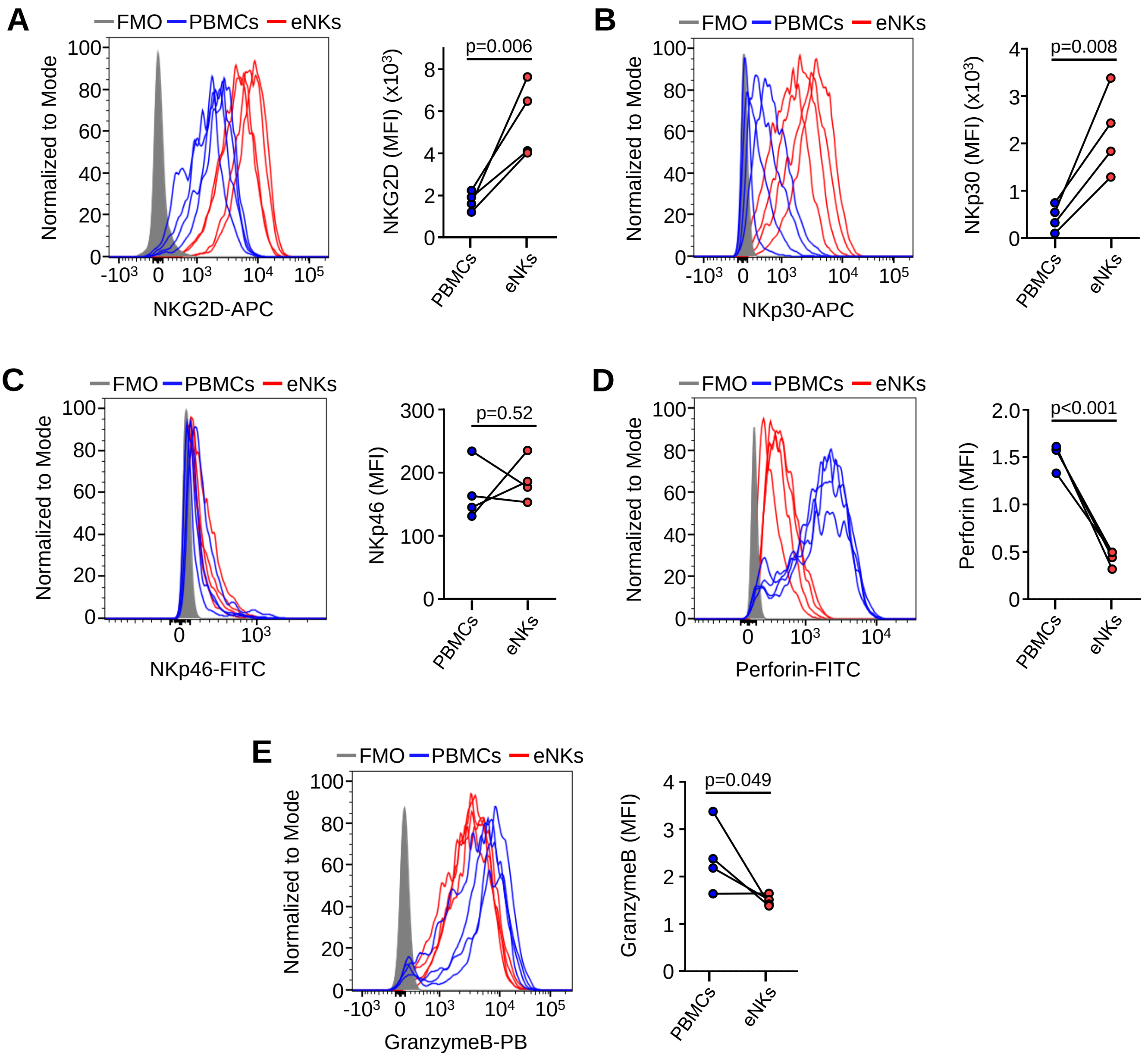


**Supplementary Figure 9. Expression of activating receptors and intracellular cytotoxic mediators by fresh and eNK cells.** eNK cells and PBMCs from four healthy blood donors were stained for surface markers including activating receptors NKG2D (**A**), NKp30 (**B**) and NKp46 (**C**) and analyzed by flow cytometry. In parallel, samples were fixed, permeabilized and stained for cytotoxic intracellular proteins Perforin (**D**), GranzymeB (**E**). Cells were gated on forward and side scatter (FSC/SSC), and NK cells were gated as CD56+ /CD3- cells.

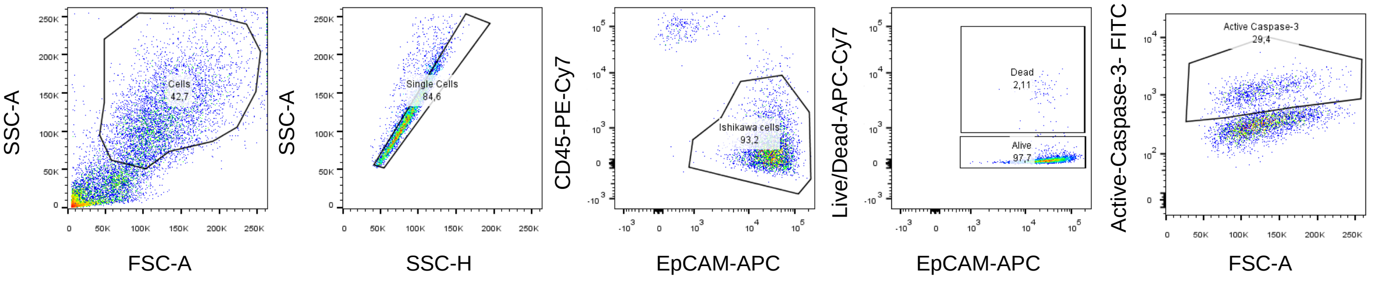


**Supplementary Figure 10. Gating strategy followed in eNK cell cytotoxicity assays.** Representative flow cytometry dot plots..

| Antibodies used in immunoblot analysis | | | | |
|---|---|---|---|---|
| **Antibody** | **Source** | **Reference** | **Dilution** | **Host** |
| TP53INP2 | In-house production | in-house | 1/400 | Mouse |
| ERK5 | Cell signaling | 3372 | 1/1000 | Rabbit |
| MEK5 | Santa Cruz | sc-365198 | 1/200 | Mouse |
| Hsp90-β | Thermo Fisher | PA3-012 | 1/16000 | Rabbit |
| FLAG | Sigma-Aldrich | F3165 | 1/100000 | Mouse |
| GST | Cell signaling | 2624 | 1/50000 | Mouse |
| Ubiquitin | Santa Cruz | sc-8017 | 1/1000 | Mouse |
| Caspase8 | BD Biosciences | 551242 | 1/400 | Mouse |
| cCaspase3 | Cell signaling | 9661 | 1/500 | Rabbit |
| β-Actin | Santa Cruz | sc-47778 | 1/5000 | Mouse |
| DR5 | Cell signaling | 8074 | 1/1000 | Rabbit |
| PARP | Cell signaling | 9542 | 1/1000 | Rabbit |
| Goat anti-Rabbit IgG (H+L) Secondary Antibody, HRP | Invitrogen | 31460 | 1/6000 | Goat |
| Goat anti-Mouse IgG (H+L) Secondary Antibody, HRP | Invitrogen | 31430 | 1/6000 | Goat |

**Supplementary Table 1**

| Antibodies used in immunofluorescent microscopy | | | | | |
|---|---|---|---|---|---|
| **Antibody** | **Conjugate** | **Source** | **Reference** | **Dilution** | **Host** |
| EpCAM | - | Cell Signalling | 2929 | 1/100 | Mouse |
| E-cadherina | - | Abcam | ab231303 | 1/500 | Mouse |
| Vimentin | - | Abcam | ab92547 | 1/250 | Rabbit |
| Phalloidin | iFluor 647 | Abcam | ab176759 | 1/100 | - |
| anti-mouse IgG | AlexaFluor 555 | Abcam | ab150118 | 1/250 | Goat |
| anti-rabbit IgG | AlexaFluor 488 | Abcam | ab150077 | 1/250 | Goat |

**Supplementary Table 2**

| Antibodies used in flow citometry | | | | | | |
|---|---|---|---|---|---|---|
| **Specificity** | **Fluorochrome** | **Clone** | **Isotype** | **Source** | **Reference** | **µl in 50 µl** |
| Human CD3 | PerCP | SK7 | mouse IgG1 | BD | 345766 | 3 |
| Human CD3 | APC-Cy7 | OKT3 | mouse IgG2a, κ | Biolegend | 317342 | 1 |
| Human CD16 | APC-eFluor 780 | CB16 | mouse IgG1, k | eBioscience | 47-0168-42 | 1 |
| Human CD45 | PE-Cy7 | HI30 | mouse IgG1, k | eBioscience | 25-0459-42 | 2 |
| Human CD56 | BV510 | NCAM16.2 | mouse IgG2b, k | BD | 563041 | 1 |
| Human CD57 | FITC | HCD57 | mouse IgM, κ | Biolegend | 322306 | 1 |
| Human CD85j (ILT2) | Pecy7 | GHI/75 | mouse IgG2b, κ | Biolegend | 333712 | 4 |
| Human CD95L (FasL) | Unconjugated | 100419 | mouse IgG2b | R&D | MAB126 | 0.1 |
| Human CD159 (NKG2A) | CFBlue | Z199 | mouse IgG2b | Dr. A. Moretta (conjugated | Not Applicable | 0.01 |
| Human CD253 (TRAIL) | PE | RIK-2 | mouse IgG1, κ | eBioscience | 12-9927-42 | 3 |
| Human CD314 (NKG2D) | APC | BAT221 | mouse IgG1 | Miltenyi Biotec | 130-092-673 | 2 |
| Human CD326 (EpCAM) | AlexaFluor-647 | 94C | mouse IgG2b, κ | Biolegend | 324212 | 0.15 |
| Human CD335 (NKp46) | FITC | 9E2 | mouse IgG1, κ | Biolegend | 331922 | 1 |
| Human CD337 (NKp30) | APC | AF29-4D12 | mouse IgG1, k | Miltenyi Biotec | 130-092-484 | 0.5 |
| **Anti-KIR mix:** | Mix of the following supernatants: | | | | | 50 |
| Human CD158a, h, g (KIR2DL1, -2DS1/S3/S5) | Hybridome supernatant | HPMA-4 | mouse IgG2b | Dr. M. López-Botet | Not Applicable | 10 |
| Human CD158b1, b2, j (KIR2DL2/L3, -2DS2) | Hybridome supernatant | CHL | mouse IgG2b | Dr. S. Ferrini | Not Applicable | 10 |
| Human CD158e1 (KIR3DL1) | Hybridome supernatant | DX9 | mouse IgG1 | Dr. L. Lanier | Not Applicable | 10 |
| Human CD158e1, k, j, i, g, e2 (KIR3DL1/L2, -2DS2/S4/S5, -3DS1) | Hybridome supernatant | 5.133 | mouse IgG1 | Dr. M. Colonna | Not Applicable | 10 |
| Human CD158f (KIR2DL5) | Hybridome supernatant | UP-R1 | mouse IgG1 | Dr. M. López-Botet | Not Applicable | 10 |
| Anti-Human active Caspase 3 | FITC | C92-605 | rabbit IgG | BD Pharmigen | 51-68654X | 3 |
| Anti-Human/mouse Granzyme B | Pacific Blue | GB11 | mouse IgG1, κ | Biolegend | 515408 | 3 |
| Anti-Human/bovine Perforin | FITC | dG9 | mouse IgG2b | BD | 556577 | 4 |
| Mouse IgG1 k isotype control | PE | P3.6.2.8.1 | mouse IgG1 k | eBioscience | 12-4714-42 | 0.3 |
| Mouse IgG2b k isotype control | PE | eBMG2b | mouse IgG2b | eBioscience | 12-4732-42 | 0.25 |
| Goat anti-mouse IgG | APC-Cy7 | polyclonal | goat | Biolegend | 405316 | 1 |
| Goat anti-Mouse IgG + IgM (H+L) | PE | polyclonal | goat | Jackson immunoResearch | 115-116-068 | 1 |

**Supplementary Table 3**

| RNA Sequencing. Differentially expressed genes (DEGs) in reponse to ERK5 inhibition | | | |
|---|---|---|---|
| EC Ishikawa cells. Log2 >\|1\|, Qvalue <0.05 | | | |
| Gene ID | Gene Symbol | log2 (ERK5i / Control) | Qvalue (ERK5i / Control) |
| 100302652 | GPR75-ASB3 | -7,627 | 0,000 |
| 100526740 | ATP5MF-PTCD1 | -6,434 | 0,000 |
| 100529144 | CORO7-PAM16 | -5,678 | 0,000 |
| 55584 | CHRNA9 | -4,821 | 0,007 |
| 144448 | TSPAN19 | -4,559 | 0,004 |
| 110091775 | C21orf59-TCP10L | -4,523 | 0,018 |
| 389396 | GLYATL3 | -4,404 | 0,022 |
| 1769 | DNAH8 | -4,284 | 0,000 |
| 10800 | CYSLTR1 | -4,281 | 0,032 |
| 6997 | TDGF1 | -4,275 | 0,040 |
| 845 | CASQ2 | -4,141 | 0,049 |
| 54084 | TSPEAR | -4,049 | 0,004 |
| 57705 | WDFY4 | -3,946 | 0,025 |
| 339761 | CYP27C1 | -3,945 | 0,024 |
| 5288 | PIK3C2G | -3,944 | 0,025 |
| 10964 | IFI44L | -3,931 | 0,000 |
| 635 | BHMT | -3,860 | 0,031 |
| 112268338 | 112268338 | -3,859 | 0,031 |
| 19 | ABCA1 | -3,720 | 0,000 |
| 441519 | CT45A3 | -3,665 | 0,000 |
| 6586 | SLIT3 | -3,604 | 0,015 |
| 94122 | SYTL5 | -3,518 | 0,000 |
| 375593 | TRIM73 | -3,485 | 0,021 |
| 27199 | OXGR1 | -3,484 | 0,021 |
| 11227 | GALNT5 | -3,407 | 0,000 |
| 8499 | PPFIA2 | -3,378 | 0,000 |
| 2565 | GABRG1 | -3,374 | 0,006 |
| 3787 | KCNS1 | -3,355 | 0,030 |
| 100509620 | LOC100509620 | -3,286 | 0,041 |
| 2568 | GABRP | -3,262 | 0,000 |
| 4693 | NDP | -3,254 | 0,000 |
| 126326 | GIPC3 | -3,213 | 0,044 |
| 245711 | SPDYA | -3,212 | 0,047 |
| 56165 | TDRD1 | -3,210 | 0,000 |
| 715 | C1R | -3,179 | 0,000 |
| 2633 | GBP1 | -3,104 | 0,000 |
| 54809 | SAMD9 | -3,030 | 0,002 |
| 146177 | VWA3A | -3,006 | 0,000 |
| 148398 | SAMD11 | -3,001 | 0,000 |
| 257177 | CFAP126 | -2,971 | 0,000 |
| 85320 | ABCC11 | -2,962 | 0,023 |
| 114801 | TMEM200A | -2,961 | 0,026 |
| 5618 | PRLR | -2,869 | 0,000 |
| 353376 | TICAM2 | -2,846 | 0,000 |
| 3910 | LAMA4 | -2,836 | 0,002 |

| | | | |
|---|---|---|---|
| 23213 | SULF1 | -2,836 | 0,038 |
| 138046 | RALYL | -2,819 | 0,000 |
| 4151 | MB | -2,818 | 0,015 |
| 285195 | SLC9A9 | -2,799 | 0,000 |
| 26266 | SLC13A4 | -2,727 | 0,010 |
| 10561 | IFI44 | -2,706 | 0,000 |
| 5104 | SERPINA5 | -2,700 | 0,049 |
| 3773 | KCNJ16 | -2,697 | 0,000 |
| 54873 | PALMD | -2,664 | 0,005 |
| 375519 | GJB7 | -2,639 | 0,013 |
| 56892 | TCIM | -2,586 | 0,000 |
| 563 | AZGP1 | -2,585 | 0,008 |
| 6323 | SCN1A | -2,547 | 0,002 |
| 3747 | KCNC2 | -2,522 | 0,000 |
| 1907 | EDN2 | -2,519 | 0,000 |
| 64078 | SLC28A3 | -2,516 | 0,000 |
| 255189 | PLA2G4F | -2,499 | 0,000 |
| 340990 | OTOG | -2,486 | 0,045 |
| 4909 | NTF4 | -2,475 | 0,008 |
| 11010 | GLIPR1 | -2,467 | 0,000 |
| 2615 | LRRC32 | -2,449 | 0,001 |
| 5788 | PTPRC | -2,448 | 0,000 |
| 4246 | SCGB2A1 | -2,435 | 0,000 |
| 57626 | KLHL1 | -2,402 | 0,000 |
| 4586 | MUC5AC | -2,394 | 0,032 |
| 10637 | LEFTY1 | -2,383 | 0,000 |
| 79962 | DNAJC22 | -2,381 | 0,000 |
| 653192 | TRIM43B | -2,377 | 0,003 |
| 101928722 | LOC101928722 | -2,375 | 0,025 |
| 6326 | SCN2A | -2,356 | 0,000 |
| 80059 | LRRTM4 | -2,355 | 0,000 |
| 283948 | NHLRC4 | -2,349 | 0,004 |
| 83876 | MRO | -2,347 | 0,000 |
| 84875 | PARP10 | -2,346 | 0,000 |
| 64005 | MYO1G | -2,341 | 0,036 |
| 26298 | EHF | -2,340 | 0,000 |
| 26154 | ABCA12 | -2,330 | 0,026 |
| 23416 | KCNH3 | -2,323 | 0,000 |
| 155465 | AGR3 | -2,322 | 0,010 |
| 124975 | GGT6 | -2,290 | 0,000 |
| 107985728 | LOC107985728 | -2,286 | 0,006 |
| 283514 | SIAH3 | -2,284 | 0,048 |
| 22989 | MYH15 | -2,283 | 0,029 |
| 8633 | UNC5C | -2,267 | 0,000 |
| 1356 | CP | -2,263 | 0,000 |
| 8111 | GPR68 | -2,243 | 0,021 |
| 55273 | TMEM100 | -2,242 | 0,024 |
| 79645 | EFCAB1 | -2,236 | 0,038 |
| 7373 | COL14A1 | -2,231 | 0,000 |
| 50853 | VILL | -2,222 | 0,000 |

| | | | |
|---|---|---|---|
| 57101 | ANO2 | -2,219 | 0,000 |
| 9619 | ABCG1 | -2,215 | 0,000 |
| 146167 | SLC38A8 | -2,211 | 0,000 |
| 84189 | SLITRK6 | -2,210 | 0,015 |
| 2678 | GGT1 | -2,195 | 0,000 |
| 94240 | EPSTI1 | -2,180 | 0,000 |
| 3488 | IGFBP5 | -2,175 | 0,000 |
| 477 | ATP1A2 | -2,173 | 0,000 |
| 374864 | CCDC178 | -2,170 | 0,000 |
| 84152 | PPP1R1B | -2,165 | 0,000 |
| 3174 | HNF4G | -2,163 | 0,000 |
| 6523 | SLC5A1 | -2,163 | 0,007 |
| 2914 | GRM4 | -2,163 | 0,000 |
| 133690 | CAPSL | -2,156 | 0,001 |
| 23452 | ANGPTL2 | -2,149 | 0,000 |
| 3778 | KCNMA1 | -2,147 | 0,000 |
| 55711 | FAR2 | -2,133 | 0,000 |
| 401190 | RGS7BP | -2,121 | 0,002 |
| 120376 | COLCA2 | -2,120 | 0,000 |
| 130813 | C2orf50 | -2,109 | 0,000 |
| 72 | ACTG2 | -2,104 | 0,000 |
| 114827 | FHAD1 | -2,094 | 0,000 |
| 129881 | CCDC173 | -2,093 | 0,000 |
| 290 | ANPEP | -2,088 | 0,000 |
| 92340 | PRR29 | -2,074 | 0,000 |
| 2001 | ELF5 | -2,067 | 0,011 |
| 158471 | PRUNE2 | -2,063 | 0,037 |
| 115111 | SLC26A7 | -2,062 | 0,000 |
| 285556 | C4orf54 | -2,061 | 0,000 |
| 260293 | CYP4X1 | -2,058 | 0,000 |
| 200162 | SPAG17 | -2,032 | 0,014 |
| 6820 | SULT2B1 | -2,030 | 0,000 |
| 399693 | CCDC187 | -2,016 | 0,004 |
| 26525 | IL36RN | -2,014 | 0,047 |
| 1281 | COL3A1 | -2,010 | 0,000 |
| 5169 | ENPP3 | -2,008 | 0,003 |
| 391723 | HELT | -2,006 | 0,036 |
| 5251 | PHEX | -2,005 | 0,035 |
| 50940 | PDE11A | -2,003 | 0,000 |
| 134121 | C5orf49 | -1,997 | 0,000 |
| 29951 | PDZRN4 | -1,992 | 0,000 |
| 284434 | NWD1 | -1,990 | 0,000 |
| 121506 | ERP27 | -1,990 | 0,007 |
| 10234 | LRRC17 | -1,986 | 0,000 |
| 827 | CAPN6 | -1,980 | 0,000 |
| 79170 | PRR15L | -1,977 | 0,000 |
| 3434 | IFIT1 | -1,972 | 0,000 |
| 55503 | TRPV6 | -1,968 | 0,000 |
| 256158 | HMCN2 | -1,964 | 0,013 |
| 2330 | FMO5 | -1,962 | 0,000 |

| | | | |
|---|---|---|---|
| 170371 | TMEM273 | -1,958 | 0,000 |
| 83872 | HMCN1 | -1,945 | 0,000 |
| 8911 | CACNA1I | -1,931 | 0,016 |
| 340061 | TMEM173 | -1,918 | 0,000 |
| 2891 | GRIA2 | -1,917 | 0,000 |
| 7044 | LEFTY2 | -1,914 | 0,005 |
| 57007 | ACKR3 | -1,905 | 0,000 |
| 344148 | NCKAP5 | -1,904 | 0,011 |
| 828 | CAPS | -1,904 | 0,000 |
| 1188 | CLCNKB | -1,899 | 0,000 |
| 1409 | CRYAA | -1,891 | 0,008 |
| 347475 | CCDC160 | -1,875 | 0,003 |
| 6565 | SLC15A2 | -1,872 | 0,000 |
| 55885 | LMO3 | -1,871 | 0,000 |
| 136306 | SVOPL | -1,862 | 0,022 |
| 94025 | MUC16 | -1,862 | 0,000 |
| 286499 | FAM133A | -1,852 | 0,004 |
| 135138 | PACRG | -1,848 | 0,016 |
| 6352 | CCL5 | -1,846 | 0,014 |
| 79740 | ZBBX | -1,837 | 0,000 |
| 29119 | CTNNA3 | -1,836 | 0,009 |
| 136288 | C7orf57 | -1,830 | 0,000 |
| 57578 | UNC79 | -1,830 | 0,000 |
| 129868 | TRIM43 | -1,822 | 0,000 |
| 64699 | TMPRSS3 | -1,813 | 0,000 |
| 2103 | ESRRB | -1,790 | 0,000 |
| 10333 | TLR6 | -1,786 | 0,025 |
| 283446 | MYO1H | -1,783 | 0,010 |
| 107984745 | LOC107984745 | -1,780 | 0,003 |
| 51313 | FAM198B | -1,778 | 0,001 |
| 222950 | NYAP1 | -1,777 | 0,000 |
| 1825 | DSC3 | -1,771 | 0,000 |
| 219681 | ARMC3 | -1,769 | 0,000 |
| 10647 | SCGB1D2 | -1,761 | 0,000 |
| 84951 | TNS4 | -1,760 | 0,000 |
| 283383 | ADGRD1 | -1,753 | 0,041 |
| 3776 | KCNK2 | -1,744 | 0,000 |
| 7481 | WNT11 | -1,744 | 0,000 |
| 5593 | PRKG2 | -1,742 | 0,000 |
| 266977 | ADGRF1 | -1,741 | 0,000 |
| 2911 | GRM1 | -1,738 | 0,000 |
| 8291 | DYSF | -1,737 | 0,000 |
| 80258 | EFHC2 | -1,731 | 0,000 |
| 642987 | TMEM232 | -1,728 | 0,023 |
| 2635 | GBP3 | -1,721 | 0,000 |
| 285533 | RNF175 | -1,721 | 0,000 |
| 978 | CDA | -1,718 | 0,000 |
| 1288 | COL4A6 | -1,704 | 0,003 |
| 56961 | SHD | -1,695 | 0,000 |
| 84000 | TMPRSS13 | -1,690 | 0,014 |

| | | | |
|---|---|---|---|
| 84889 | SLC7A3 | -1,689 | 0,030 |
| 5349 | FXYD3 | -1,685 | 0,000 |
| 9630 | GNA14 | -1,674 | 0,005 |
| 283078 | MKX | -1,671 | 0,000 |
| 55 | ACPP | -1,661 | 0,000 |
| 4778 | NFE2 | -1,652 | 0,000 |
| 341019 | DCDC1 | -1,651 | 0,000 |
| 200931 | SLC51A | -1,650 | 0,000 |
| 389434 | IYD | -1,647 | 0,027 |
| 254050 | LRRC43 | -1,640 | 0,007 |
| 56144 | PCDHA4 | -1,638 | 0,000 |
| 56138 | PCDHA11 | -1,635 | 0,000 |
| 25878 | MXRA5 | -1,633 | 0,000 |
| 952 | CD38 | -1,617 | 0,000 |
| 140803 | TRPM6 | -1,616 | 0,002 |
| 2581 | GALC | -1,604 | 0,000 |
| 4057 | LTF | -1,603 | 0,020 |
| 11118 | BTN3A2 | -1,602 | 0,000 |
| 7222 | TRPC3 | -1,599 | 0,000 |
| 117283 | IP6K3 | -1,591 | 0,000 |
| 9623 | TCL1B | -1,587 | 0,000 |
| 6920 | TCEA3 | -1,585 | 0,000 |
| 10551 | AGR2 | -1,577 | 0,000 |
| 79630 | C1orf54 | -1,567 | 0,006 |
| 11155 | LDB3 | -1,567 | 0,008 |
| 285386 | TPRG1 | -1,561 | 0,000 |
| 50507 | NOX4 | -1,561 | 0,000 |
| 718 | C3 | -1,556 | 0,000 |
| 8483 | CILP | -1,552 | 0,000 |
| 56649 | TMPRSS4 | -1,551 | 0,000 |
| 91584 | PLXNA4 | -1,549 | 0,026 |
| 338811 | FAM19A2 | -1,548 | 0,009 |
| 83449 | PMFBP1 | -1,547 | 0,031 |
| 5348 | FXYD1 | -1,547 | 0,002 |
| 79919 | MAB21L4 | -1,547 | 0,000 |
| 167410 | LIX1 | -1,532 | 0,001 |
| 286464 | CFAP47 | -1,532 | 0,004 |
| 93082 | NEURL3 | -1,531 | 0,002 |
| 1475 | CSTA | -1,526 | 0,013 |
| 55806 | HR | -1,520 | 0,000 |
| 79370 | BCL2L14 | -1,519 | 0,031 |
| 56477 | CCL28 | -1,518 | 0,000 |
| 3625 | INHBB | -1,516 | 0,046 |
| 8743 | TNFSF10 | -1,510 | 0,000 |
| 25876 | SPEF1 | -1,505 | 0,002 |
| 58494 | JAM2 | -1,504 | 0,000 |
| 89870 | TRIM15 | -1,497 | 0,023 |
| 51702 | PADI3 | -1,490 | 0,000 |
| 80270 | HSD3B7 | -1,489 | 0,000 |
| 368 | ABCC6 | -1,481 | 0,000 |

| | | | |
|---|---|---|---|
| 4330 | MN1 | -1,478 | 0,005 |
| 251 | ALPG | -1,477 | 0,000 |
| 5241 | PGR | -1,471 | 0,000 |
| 8519 | IFITM1 | -1,469 | 0,000 |
| 441869 | ANKRD65 | -1,459 | 0,001 |
| 146802 | SLC47A2 | -1,447 | 0,000 |
| 2697 | GJA1 | -1,440 | 0,000 |
| 144535 | CFAP54 | -1,439 | 0,000 |
| 3081 | HGD | -1,436 | 0,000 |
| 844 | CASQ1 | -1,426 | 0,000 |
| 201181 | ZNF385C | -1,419 | 0,003 |
| 25840 | METTL7A | -1,419 | 0,000 |
| 116535 | MRGPRF | -1,418 | 0,000 |
| 50632 | CALY | -1,415 | 0,000 |
| 200373 | CFAP221 | -1,414 | 0,000 |
| 26585 | GREM1 | -1,413 | 0,008 |
| 8742 | TNFSF12 | -1,413 | 0,000 |
| 51050 | PI15 | -1,407 | 0,000 |
| 79839 | CCDC102B | -1,404 | 0,018 |
| 353345 | GPR141 | -1,402 | 0,015 |
| 57158 | JPH2 | -1,400 | 0,007 |
| 1767 | DNAH5 | -1,390 | 0,002 |
| 56898 | BDH2 | -1,389 | 0,000 |
| 8549 | LGR5 | -1,389 | 0,000 |
| 57639 | CCDC146 | -1,383 | 0,000 |
| 9068 | ANGPTL1 | -1,381 | 0,000 |
| 59 | ACTA2 | -1,374 | 0,000 |
| 398 | ARHGDIG | -1,370 | 0,000 |
| 728464 | METTL24 | -1,370 | 0,024 |
| 29116 | MYLIP | -1,369 | 0,000 |
| 199920 | FYB2 | -1,367 | 0,000 |
| 5454 | POU3F2 | -1,363 | 0,037 |
| 101060179 | LOC101060179 | -1,356 | 0,000 |
| 101928764 | LOC101928764 | -1,355 | 0,008 |
| 374907 | B3GNT8 | -1,355 | 0,010 |
| 56925 | LXN | -1,353 | 0,000 |
| 2444 | FRK | -1,351 | 0,000 |
| 81578 | COL21A1 | -1,351 | 0,001 |
| 2348 | FOLR1 | -1,350 | 0,000 |
| 79858 | NEK11 | -1,346 | 0,000 |
| 57514 | ARHGAP31 | -1,346 | 0,000 |
| 55691 | FRMD4A | -1,344 | 0,000 |
| 5896 | RAG1 | -1,341 | 0,000 |
| 8382 | NME5 | -1,341 | 0,002 |
| 148418 | SAMD13 | -1,341 | 0,000 |
| 5176 | SERPINF1 | -1,338 | 0,000 |
| 128153 | SPATA17 | -1,337 | 0,000 |
| 79983 | POF1B | -1,336 | 0,005 |
| 133688 | UGT3A1 | -1,336 | 0,034 |
| 200810 | ALG1L | -1,335 | 0,007 |

| | | | |
|---|---|---|---|
| 4810 | NHS | -1,334 | 0,000 |
| 79838 | TMC5 | -1,332 | 0,000 |
| 54436 | SH3TC1 | -1,330 | 0,000 |
| 64170 | CARD9 | -1,326 | 0,028 |
| 83468 | GLT8D2 | -1,321 | 0,000 |
| 57758 | SCUBE2 | -1,315 | 0,000 |
| 9077 | DIRAS3 | -1,311 | 0,012 |
| 770 | CA11 | -1,311 | 0,000 |
| 349667 | RTN4RL2 | -1,310 | 0,010 |
| 118932 | ANKRD22 | -1,310 | 0,000 |
| 287 | ANK2 | -1,305 | 0,000 |
| 25790 | CFAP45 | -1,304 | 0,000 |
| 1264 | CNN1 | -1,302 | 0,000 |
| 54103 | GSAP | -1,298 | 0,000 |
| 10804 | GJB6 | -1,297 | 0,044 |
| 2566 | GABRG2 | -1,297 | 0,008 |
| 147138 | TMC8 | -1,293 | 0,000 |
| 6862 | TBXT | -1,293 | 0,016 |
| 57664 | PLEKHA4 | -1,293 | 0,000 |
| 7539 | ZFP37 | -1,292 | 0,000 |
| 91624 | NEXN | -1,291 | 0,000 |
| 151516 | ASPRV1 | -1,290 | 0,001 |
| 3990 | LIPC | -1,288 | 0,001 |
| 113 | ADCY7 | -1,288 | 0,000 |
| 9256 | TSPOAP1 | -1,288 | 0,000 |
| 401546 | C9orf152 | -1,285 | 0,028 |
| 149840 | SHLD1 | -1,282 | 0,000 |
| 3547 | IGSF1 | -1,281 | 0,000 |
| 168002 | DACT2 | -1,274 | 0,000 |
| 345895 | RSPH4A | -1,272 | 0,000 |
| 23632 | CA14 | -1,272 | 0,034 |
| 494514 | TYMSOS | -1,262 | 0,000 |
| 4143 | MAT1A | -1,260 | 0,042 |
| 148170 | CDC42EP5 | -1,259 | 0,044 |
| 80217 | CFAP43 | -1,259 | 0,000 |
| 54933 | RHBDL2 | -1,258 | 0,000 |
| 23639 | LRRC6 | -1,257 | 0,000 |
| 23345 | SYNE1 | -1,257 | 0,000 |
| 90557 | CCDC74A | -1,250 | 0,000 |
| 7036 | TFR2 | -1,250 | 0,015 |
| 55601 | DDX60 | -1,249 | 0,000 |
| 9283 | GPR37L1 | -1,249 | 0,038 |
| 2823 | GPM6A | -1,247 | 0,002 |
| 3242 | HPD | -1,246 | 0,004 |
| 152110 | NEK10 | -1,242 | 0,000 |
| 55036 | CCDC40 | -1,240 | 0,000 |
| 141 | ADPRH | -1,239 | 0,000 |
| 146754 | DNAH2 | -1,238 | 0,003 |
| 79656 | BEND5 | -1,238 | 0,000 |
| 105373926 | 105373926 | -1,237 | 0,000 |

| | | | |
|---|---|---|---|
| 51062 | ATL1 | -1,235 | 0,000 |
| 51364 | ZMYND10 | -1,235 | 0,000 |
| 27285 | TEKT2 | -1,234 | 0,000 |
| 25928 | SOSTDC1 | -1,232 | 0,000 |
| 397 | ARHGDIB | -1,227 | 0,000 |
| 9886 | RHOBTB1 | -1,226 | 0,000 |
| 400986 | ANKRD36C | -1,223 | 0,000 |
| 3852 | KRT5 | -1,222 | 0,000 |
| 7851 | MALL | -1,220 | 0,000 |
| 4118 | MAL | -1,220 | 0,000 |
| 3487 | IGFBP4 | -1,218 | 0,000 |
| 259232 | NALCN | -1,215 | 0,000 |
| 1462 | VCAN | -1,212 | 0,000 |
| 822 | CAPG | -1,210 | 0,000 |
| 8643 | PTCH2 | -1,203 | 0,000 |
| 159989 | DEUP1 | -1,198 | 0,028 |
| 57125 | PLXDC1 | -1,198 | 0,029 |
| 1299 | COL9A3 | -1,192 | 0,000 |
| 8578 | SCARF1 | -1,191 | 0,034 |
| 79814 | AGMAT | -1,190 | 0,000 |
| 22987 | SV2C | -1,189 | 0,000 |
| 56164 | STK31 | -1,188 | 0,000 |
| 10083 | USH1C | -1,186 | 0,018 |
| 57161 | PELI2 | -1,181 | 0,000 |
| 56171 | DNAH7 | -1,179 | 0,000 |
| 80731 | THSD7B | -1,177 | 0,000 |
| 51131 | PHF11 | -1,175 | 0,000 |
| 4313 | MMP2 | -1,173 | 0,000 |
| 3433 | IFIT2 | -1,170 | 0,000 |
| 558 | AXL | -1,166 | 0,000 |
| 2099 | ESR1 | -1,165 | 0,000 |
| 5139 | PDE3A | -1,164 | 0,000 |
| 101059918 | GOLGA8R | -1,162 | 0,000 |
| 22874 | PLEKHA6 | -1,161 | 0,000 |
| 6899 | TBX1 | -1,159 | 0,015 |
| 54716 | SLC6A20 | -1,159 | 0,000 |
| 140469 | MYO3B | -1,156 | 0,000 |
| 8082 | SSPN | -1,154 | 0,000 |
| 11248 | NXPH3 | -1,148 | 0,000 |
| 55205 | ZNF532 | -1,148 | 0,000 |
| 57214 | CEMIP | -1,144 | 0,003 |
| 9625 | AATK | -1,141 | 0,000 |
| 55679 | LIMS2 | -1,141 | 0,000 |
| 3232 | HOXD3 | -1,141 | 0,022 |
| 3431 | SP110 | -1,140 | 0,014 |
| 342979 | PALM3 | -1,139 | 0,000 |
| 286 | ANK1 | -1,136 | 0,013 |
| 388115 | CCDC9B | -1,135 | 0,000 |
| 79628 | SH3TC2 | -1,133 | 0,030 |
| 6414 | SELENOP | -1,131 | 0,000 |

| | | | |
|---|---|---|---|
| 135228 | CD109 | -1,129 | 0,000 |
| 215 | ABCD1 | -1,122 | 0,000 |
| 223117 | SEMA3D | -1,119 | 0,000 |
| 117581 | TWIST2 | -1,117 | 0,007 |
| 3784 | KCNQ1 | -1,115 | 0,000 |
| 3754 | KCNF1 | -1,114 | 0,000 |
| 23150 | FRMD4B | -1,112 | 0,000 |
| 7043 | TGFB3 | -1,110 | 0,000 |
| 1016 | CDH18 | -1,109 | 0,000 |
| 139411 | PTCHD1 | -1,109 | 0,000 |
| 28968 | SLC6A16 | -1,107 | 0,028 |
| 57232 | ZNF630 | -1,105 | 0,000 |
| 479 | ATP12A | -1,105 | 0,017 |
| 112495 | GTF3C6 | -1,105 | 0,000 |
| 6536 | SLC6A9 | -1,104 | 0,000 |
| 54626 | HES2 | -1,104 | 0,000 |
| 120892 | LRRK2 | -1,102 | 0,000 |
| 80763 | SPX | -1,101 | 0,000 |
| 22900 | CARD8 | -1,098 | 0,000 |
| 11119 | BTN3A1 | -1,095 | 0,000 |
| 195814 | SDR16C5 | -1,092 | 0,032 |
| 728047 | GOLGA8O | -1,089 | 0,000 |
| 90427 | BMF | -1,088 | 0,000 |
| 6567 | SLC16A2 | -1,087 | 0,000 |
| 64798 | DEPTOR | -1,084 | 0,000 |
| 56901 | NDUFA4L2 | -1,081 | 0,000 |
| 29800 | ZDHHC1 | -1,080 | 0,000 |
| 51302 | CYP39A1 | -1,078 | 0,000 |
| 5645 | PRSS2 | -1,075 | 0,000 |
| 79819 | WDR78 | -1,073 | 0,000 |
| 552 | AVPR1A | -1,072 | 0,006 |
| 10267 | RAMP1 | -1,071 | 0,000 |
| 7113 | TMPRSS2 | -1,068 | 0,000 |
| 5027 | P2RX7 | -1,065 | 0,039 |
| 255394 | TCP11L2 | -1,064 | 0,000 |
| 5118 | PCOLCE | -1,064 | 0,000 |
| 158866 | ZDHHC15 | -1,063 | 0,002 |
| 85452 | CFAP74 | -1,061 | 0,003 |
| 2651 | GCNT2 | -1,061 | 0,000 |
| 10581 | IFITM2 | -1,060 | 0,000 |
| 23236 | PLCB1 | -1,056 | 0,000 |
| 8622 | PDE8B | -1,055 | 0,000 |
| 9435 | CHST2 | -1,054 | 0,000 |
| 56956 | LHX9 | -1,052 | 0,001 |
| 2104 | ESRRG | -1,050 | 0,011 |
| 79815 | NIPAL2 | -1,050 | 0,000 |
| 30818 | KCNIP3 | -1,049 | 0,000 |
| 400746 | NCMAP | -1,049 | 0,000 |
| 112267992 | LOC112267992 | -1,045 | 0,000 |
| 55315 | SLC29A3 | -1,044 | 0,000 |

| | | | |
|---|---|---|---|
| 745 | MYRF | -1,043 | 0,000 |
| 54768 | HYDIN | -1,040 | 0,000 |
| 342850 | ANKRD62 | -1,037 | 0,015 |
| 80319 | CXXC4 | -1,035 | 0,003 |
| 11174 | ADAMTS6 | -1,035 | 0,011 |
| 56105 | PCDHGA11 | -1,034 | 0,000 |
| 10319 | LAMC3 | -1,032 | 0,000 |
| 128344 | PIFO | -1,030 | 0,000 |
| 63970 | TP53AIP1 | -1,030 | 0,002 |
| 6445 | SGCG | -1,030 | 0,006 |
| 3233 | HOXD4 | -1,029 | 0,000 |
| 132014 | IL17RE | -1,027 | 0,000 |
| 150696 | PROM2 | -1,026 | 0,000 |
| 100129583 | FAM47E | -1,025 | 0,002 |
| 343990 | KIAA1211L | -1,025 | 0,000 |
| 1382 | CRABP2 | -1,025 | 0,000 |
| 154091 | SLC2A12 | -1,024 | 0,000 |
| 89796 | NAV1 | -1,024 | 0,000 |
| 56098 | PCDHGC4 | -1,022 | 0,000 |
| 6296 | ACSM3 | -1,021 | 0,000 |
| 24 | ABCA4 | -1,017 | 0,000 |
| 51686 | OAZ3 | -1,015 | 0,045 |
| 9454 | HOMER3 | -1,014 | 0,000 |
| 2346 | FOLH1 | -1,014 | 0,000 |
| 94120 | SYTL3 | -1,011 | 0,000 |
| 8828 | NRP2 | -1,009 | 0,032 |
| 126231 | ZNF573 | -1,009 | 0,000 |
| 1278 | COL1A2 | -1,009 | 0,000 |
| 57020 | VPS35L | -1,008 | 0,000 |
| 171019 | ADAMTS19 | -1,007 | 0,000 |
| 56108 | PCDHGA7 | -1,007 | 0,000 |
| 56131 | PCDHB4 | -1,005 | 0,035 |
| 343450 | KCNT2 | -1,004 | 0,023 |
| 2256 | FGF11 | -1,001 | 0,000 |
| 9982 | FGFBP1 | 1,002 | 0,000 |
| 79639 | TMEM53 | 1,003 | 0,000 |
| 283208 | P4HA3 | 1,003 | 0,000 |
| 39 | ACAT2 | 1,005 | 0,000 |
| 6812 | STXBP1 | 1,006 | 0,000 |
| 221491 | SMIM29 | 1,012 | 0,000 |
| 7108 | TM7SF2 | 1,017 | 0,000 |
| 57493 | HEG1 | 1,017 | 0,000 |
| 57134 | MAN1C1 | 1,020 | 0,000 |
| 6307 | MSMO1 | 1,021 | 0,000 |
| 79825 | EFCC1 | 1,025 | 0,014 |
| 130399 | ACVR1C | 1,029 | 0,000 |
| 22979 | EFR3B | 1,031 | 0,000 |
| 388135 | INSYN1 | 1,038 | 0,031 |
| 116412 | ZNF837 | 1,038 | 0,000 |
| 54769 | DIRAS2 | 1,038 | 0,024 |

| | | | |
|---|---|---|---|
| 653082 | ZDHHC11B | 1,038 | 0,000 |
| 1113 | CHGA | 1,043 | 0,000 |
| 80339 | PNPLA3 | 1,045 | 0,000 |
| 644596 | SMIM10L2B | 1,056 | 0,000 |
| 9052 | GPRC5A | 1,057 | 0,000 |
| 7111 | TMOD1 | 1,064 | 0,000 |
| 348262 | MCRIP1 | 1,067 | 0,000 |
| 3992 | FADS1 | 1,068 | 0,000 |
| 115123 | MARCH3 | 1,069 | 0,000 |
| 3665 | IRF7 | 1,077 | 0,000 |
| 2264 | FGFR4 | 1,077 | 0,000 |
| 3006 | HIST1H1C | 1,077 | 0,000 |
| 2026 | ENO2 | 1,080 | 0,000 |
| 8349 | HIST2H2BE | 1,081 | 0,000 |
| 9645 | MICAL2 | 1,085 | 0,000 |
| 9066 | SYT7 | 1,090 | 0,000 |
| 8120 | AP3B2 | 1,092 | 0,000 |
| 2224 | FDPS | 1,096 | 0,000 |
| 51557 | LGSN | 1,098 | 0,017 |
| 10411 | RAPGEF3 | 1,105 | 0,000 |
| 151176 | ERFE | 1,108 | 0,000 |
| 651746 | ANKRD33B | 1,113 | 0,000 |
| 79629 | OCEL1 | 1,113 | 0,000 |
| 1848 | DUSP6 | 1,115 | 0,000 |
| 47 | ACLY | 1,118 | 0,000 |
| 84978 | FRMD5 | 1,120 | 0,000 |
| 273 | AMPH | 1,121 | 0,000 |
| 353189 | SLCO4C1 | 1,121 | 0,000 |
| 3748 | KCNC3 | 1,123 | 0,000 |
| 28231 | SLCO4A1 | 1,123 | 0,000 |
| 51181 | DCXR | 1,129 | 0,000 |
| 285966 | TCAF2 | 1,131 | 0,019 |
| 26575 | RGS17 | 1,133 | 0,040 |
| 92840 | REEP6 | 1,135 | 0,000 |
| 92558 | BICDL1 | 1,137 | 0,000 |
| 104 | ADARB1 | 1,146 | 0,000 |
| 5270 | SERPINE2 | 1,147 | 0,000 |
| 90113 | VWA5B2 | 1,148 | 0,000 |
| 2034 | EPAS1 | 1,149 | 0,000 |
| 1030 | CDKN2B | 1,150 | 0,000 |
| 6456 | SH3GL2 | 1,157 | 0,000 |
| 6853 | SYN1 | 1,158 | 0,000 |
| 3157 | HMGCS1 | 1,163 | 0,000 |
| 3949 | LDLR | 1,177 | 0,000 |
| 8347 | HIST1H2BC | 1,178 | 0,000 |
| 23175 | LPIN1 | 1,181 | 0,000 |
| 2118 | ETV4 | 1,184 | 0,000 |
| 4277 | MICB | 1,206 | 0,000 |
| 143425 | SYT9 | 1,207 | 0,013 |
| 3632 | INPP5A | 1,222 | 0,000 |

| | | | |
|---|---|---|---|
| 56155 | TEX14 | 1,227 | 0,000 |
| 113177 | IZUMO4 | 1,233 | 0,025 |
| 162494 | RHBDL3 | 1,238 | 0,000 |
| 374 | AREG | 1,239 | 0,003 |
| 10580 | SORBS1 | 1,240 | 0,000 |
| 8326 | FZD9 | 1,242 | 0,000 |
| 8553 | BHLHE40 | 1,245 | 0,000 |
| 64129 | TINAGL1 | 1,250 | 0,000 |
| 8970 | HIST1H2BJ | 1,250 | 0,000 |
| 4744 | NEFH | 1,251 | 0,000 |
| 84953 | MICALCL | 1,252 | 0,000 |
| 8927 | BSN | 1,259 | 0,000 |
| 80726 | IQCN | 1,269 | 0,000 |
| 5979 | RET | 1,280 | 0,000 |
| 100507436 | MICA | 1,286 | 0,000 |
| 55512 | SMPD3 | 1,291 | 0,007 |
| 9518 | GDF15 | 1,298 | 0,000 |
| 84915 | FAM222A | 1,299 | 0,000 |
| 4884 | NPTX1 | 1,300 | 0,019 |
| 27019 | DNAI1 | 1,300 | 0,010 |
| 8969 | HIST1H2AG | 1,300 | 0,000 |
| 51421 | AMOTL2 | 1,302 | 0,000 |
| 81848 | SPRY4 | 1,308 | 0,000 |
| 55902 | ACSS2 | 1,315 | 0,000 |
| 8909 | ENDOU | 1,315 | 0,000 |
| 1969 | EPHA2 | 1,322 | 0,000 |
| 79012 | CAMKV | 1,332 | 0,040 |
| 55502 | HES6 | 1,350 | 0,000 |
| 23764 | MAFF | 1,359 | 0,004 |
| 6457 | SH3GL3 | 1,366 | 0,000 |
| 3976 | LIF | 1,366 | 0,000 |
| 7093 | TLL2 | 1,377 | 0,000 |
| 81788 | NUAK2 | 1,382 | 0,000 |
| 389692 | MAFA | 1,387 | 0,000 |
| 8651 | SOCS1 | 1,388 | 0,000 |
| 2194 | FASN | 1,397 | 0,000 |
| 7425 | VGF | 1,405 | 0,043 |
| 1846 | DUSP4 | 1,422 | 0,000 |
| 9586 | CREB5 | 1,443 | 0,003 |
| 8358 | HIST1H3B | 1,443 | 0,037 |
| 3084 | NRG1 | 1,457 | 0,000 |
| 8395 | PIP5K1B | 1,459 | 0,000 |
| 54855 | TENT5C | 1,465 | 0,000 |
| 1839 | HBEGF | 1,467 | 0,000 |
| 201625 | DNAH12 | 1,468 | 0,014 |
| 81544 | GDPD5 | 1,479 | 0,000 |
| 4880 | NPPC | 1,486 | 0,003 |
| 85409 | NKD2 | 1,488 | 0,000 |
| 51440 | HPCAL4 | 1,489 | 0,000 |
| 54567 | DLL4 | 1,504 | 0,016 |

| | | | |
|---|---|---|---|
| 10202 | DHRS2 | 1,506 | 0,000 |
| 392255 | GDF6 | 1,513 | 0,028 |
| 54361 | WNT4 | 1,529 | 0,016 |
| 7980 | TFPI2 | 1,537 | 0,000 |
| 57468 | SLC12A5 | 1,546 | 0,021 |
| 4047 | LSS | 1,554 | 0,000 |
| 143282 | FGFBP3 | 1,582 | 0,000 |
| 55652 | SLC48A1 | 1,588 | 0,000 |
| 3422 | IDI1 | 1,591 | 0,000 |
| 23237 | ARC | 1,616 | 0,000 |
| 6425 | SFRP5 | 1,621 | 0,005 |
| 57718 | PPP4R4 | 1,640 | 0,000 |
| 654790 | PCP4L1 | 1,640 | 0,000 |
| 387763 | C11orf96 | 1,648 | 0,008 |
| 124976 | SPNS2 | 1,655 | 0,001 |
| 4584 | MUC3A | 1,668 | 0,010 |
| 9248 | GPR50 | 1,675 | 0,000 |
| 4597 | MVD | 1,732 | 0,000 |
| 716 | C1S | 1,734 | 0,000 |
| 51513 | ETV7 | 1,742 | 0,000 |
| 8365 | HIST1H4H | 1,770 | 0,000 |
| 10570 | DPYSL4 | 1,799 | 0,005 |
| 57624 | NYAP2 | 1,803 | 0,046 |
| 161725 | OTUD7A | 1,817 | 0,000 |
| 9001 | HAP1 | 1,826 | 0,000 |
| 3638 | INSIG1 | 1,829 | 0,000 |
| 230 | ALDOC | 1,853 | 0,000 |
| 729877 | TBC1D3H | 1,913 | 0,041 |
| 3791 | KDR | 1,935 | 0,017 |
| 55824 | PAG1 | 1,944 | 0,000 |
| 81872 | KRTAP2-1 | 1,948 | 0,000 |
| 11076 | TPPP | 1,963 | 0,000 |
| 114818 | KLHL29 | 1,969 | 0,000 |
| 25907 | TMEM158 | 1,975 | 0,000 |
| 102724862 | TBC1D3I | 2,002 | 0,033 |
| 128312 | HIST3H2BB | 2,014 | 0,002 |
| 730755 | KRTAP2-3 | 2,018 | 0,000 |
| 55567 | DNAH3 | 2,072 | 0,000 |
| 1809 | DPYSL3 | 2,105 | 0,000 |
| 345611 | IRGM | 2,115 | 0,016 |
| 6752 | SSTR2 | 2,140 | 0,002 |
| 6620 | SNCB | 2,245 | 0,000 |
| 23205 | ACSBG1 | 2,249 | 0,000 |
| 50615 | IL21R | 2,251 | 0,004 |
| 654 | BMP6 | 2,256 | 0,000 |
| 869 | CBLN1 | 2,360 | 0,004 |
| 7634 | ZNF80 | 2,377 | 0,000 |
| 8626 | TP63 | 2,468 | 0,000 |
| 4885 | NPTX2 | 2,523 | 0,000 |
| 115362 | GBP5 | 2,600 | 0,000 |

| | | | |
|---|---|---|---|
| 1233 | CCR4 | 2,606 | 0,000 |
| 4804 | NGFR | 2,624 | 0,000 |
| 7781 | SLC30A3 | 2,921 | 0,000 |
| 125931 | CEACAM20 | 3,014 | 0,000 |
| 125965 | COX6B2 | 3,039 | 0,020 |
| 360200 | TMPRSS9 | 3,160 | 0,000 |
| 51458 | RHCG | 3,207 | 0,043 |
| 5453 | POU3F1 | 3,500 | 0,022 |
| 6123 | RPL3L | 4,133 | 0,000 |
| 8529 | CYP4F2 | 4,256 | 0,012 |
| 63976 | PRDM16 | 4,357 | 0,031 |
| 8789 | FBP2 | 4,452 | 0,000 |
